# In-cell discovery and characterization of a non-canonical bacterial protein translocation-folding complex

**DOI:** 10.1101/2025.04.25.650208

**Authors:** Rasmus K. Jensen, Liang Xue, Federico Marotta, Joseph C. Somody, Joel Selkrig, Swantje Lenz, Juri Rappsilber, Mikhail M. Savitski, Jan Kosinski, Athanasios Typas, Maria Zimmermann-Kogadeeva, Peer Bork, Julia Mahamid

**Affiliations:** Molecular Systems Biology Unit, European Molecular Biology Laboratory (EMBL), Meyerhofstraße 1, 69117 Heidelberg, Germany; Current address: Key Laboratory of Biomacromolecules, Institute of Biophysics, Chinese Academy of Sciences, Beijing 100101, China; Julius-Maximilians-Universität Würzburg, Fakultät für Biologie, Am Hubland, 97074 Würzburg, Germany; Genome Biology Unit, EMBL, Meyerhofstraße 1, 69117 Heidelberg, Germany; Institute of Medical Microbiology, RWTH University Hospital, Aachen 52074, Germany; Bioanalytics Unit, Institute of Biotechnology, Technische Universität Berlin, 10623 Berlin, Germany; Max Planck Institute of Molecular Cell Biology and Genetics, Pfotenhauerstraße 1 08, 01307 Dresden, Germany; EMBL, Hamburg, Notkestrasse 85, 20607 Hamburg, Germany; Centre of Structural Systems Biology, Notkestrasse 85, 20607 Hamburg, Germany; Cell biology and Biophysics Unit, EMBL, Meyerhofstraße 1, 69117 Heidelberg, Germany

## Abstract

Cryo-electron tomography has emerged as powerful technology for in-cell structural biology, and in combination with breakthroughs in protein structure prediction, offers a unique opportunity for illuminating functions of previously uncharacterized macromolecular complexes. Here, we used the genome-reduced bacterium *Mycoplasma pneumoniae* as a minimal cell model, and determined in-cell maps of an unknown complex located at the cell surface. By combining proteomics, structure prediction, and bioinformatics, we identified the complex to include the conserved Sec-translocon and an extracellular dome-like structure largely formed by three uncharacterized proteins (Mdps). The Mdps show structural homology to a periplasmic ATP-independent foldase, PrsA, with Mdp444 retaining key catalytic residues. Our study provides the first sub-nanometer resolution maps of the complete bacterial Sec-translocation machinery, depicting several conformational states that allow us to suggest a model for translocation initiation, and revealing its coordination with a novel, extracellular, protein-folding system.

## Introduction

Cryogenic electron tomography (cryo-ET) has revolutionized in-cell structural biology, enabling mechanistic studies of macromolecular complexes in their native environment^1,2^. Recent efforts in optimizing sample preparation^3–6^, data acquisition strategies^7–9^, and downstream data processing^10,11^ allow large, high-resolution, datasets in intact cells to be acquired and analyzed. As larger volumes of data and advanced analysis tools become available, cryo-ET can start addressing lower abundance and more diverse macromolecular species. However, assigning identity and biological function to structures observed *in situ* across diverse organisms remains a challenge, especially given that many complexes have not been previously characterized by classical *in vitro* structural biology methods, and can typically only be resolved to intermediate resolution *in situ*^12–14^. In this study, we address such a challenge using the bacterium *Mycoplasma pneumoniae*. This genome-reduced human pathogen is one of the simplest self-replicating organisms, and has been extensively used as a minimal cell model for systems biology^15,16^. It therefore benefits from a broad range of available data resources, including whole-cell crosslinking mass spectrometry^17^, protein abundance measurements^18^, and whole-genome gene-essentiality studies^19^. *M. pneumoniae* has also proved a suitable model for in-cell cryo-ET studies, as it naturally adheres to substrates and can thus be grown directly on cryo-electron microscopy (cryo-EM) grids, and its small cell size enables imaging without the requirement of sample thinning^17^. This allows for the acquisition of large-scale, high-quality cryo-ET datasets, facilitating high-resolution in-cell structure determination, as demonstrated for ribosomes resolved to 3.5 Å in native cells^20^. Despite its minimal genome, containing less than 700 protein-coding sequences and its widespread use as a minimal cell model, over 20% of the *M. pneumoniae* proteome remains uncharacterized^21,22^, making it an ideal candidate for *de novo* structural discovery with cryo-ET.

Here, we focused on the characterization of a previously unknown membrane-bound protein complex visualized on the *M. pneumoniae* cell surface. Through a combination of proteomics, structure prediction and integrative structural biology tools, we determined its molecular composition. Several of the proteins were of unknown function, and we harnessed structural homology search tools, phylogenetic analysis and cellular perturbation experiments to elucidate that the extracellular proteins formed a protein-folding chamber at the cell surface. Strikingly, we found this novel complex to constitutively associate with the Sec-translocation machinery, suggesting a functional link between the two systems.

Protein translocation across the membrane is a fundamental process in all living cells, and in *M. pneumoniae*, the Sec machinery represents the only known translocation or secretion system^23,24^. The Sec translocation complex is evolutionarily conserved across bacteria, archaea, and eukaryotes, where it facilitates protein translocation and membrane protein insertion. In bacteria, the central translocation pore is formed by the three transmembrane proteins SecY, SecE and SecG, and is often denoted SecYEG^25^. The SecYEG channel further associates with SecD, SecF, YidC and YajC forming a holotranslocon^26–28^. SecD and SecF are necessary for efficient translocation *in vivo*^29^, and have fused into a single protein, SecDF, in *Mycoplasma*^24^. Although its precise mechanism remains unclear, SecDF is thought to harness the proton motive force to pull translocated substrates at the extracellular side of the cell membrane^29–31^. SecA is an intracellular ATPase, which binds newly translated proteins (preprotein) and recruit them to the Sec-translocon. SecA is mainly known to bind preproteins post-translationally, but can also act co-translationally for some proteins^32–34^. After association of preprotein-bound SecA with SecYEG, SecA drives translocation through ATP hydrolysis^34^. SecA is also important for maintaining the preprotein in the correct state for translocation, as the Sec machinery mainly translocates unfolded proteins^35^. Although many structures of individual Sec components and the SecA-SecY complex exist, the structural and functional organization of the holotranslocon remains unclear^28,31,36–39^. In its native environment of the bacterial membrane, the holotranslocon is expected to be largely fully membrane-embedded, making it a challenging target for in-cell structural studies. Here, guided by its association with the novel extracellular dome complex, we resolved a 9.1 Å in-cell map of the holotranslocon, unveiling its architecture and interaction network within the bacterial membrane. Our detailed classification scheme revealed distinct conformational states, providing insights into the molecular mechanism of bacterial translocation initiation.

## Results

### In-cell architecture of a dome-like transmembrane complex in *M. pneumoniae*

Cryo-ET data of *M. pneumoniae* cells revealed the presence of a large membrane-associated, dome-like structure extending into the extracellular space (Figure 1A, Methods). We detected 39 ± 17 complexes per cell (Figure 1B), with 38% of the complexes forming clusters, typically in groups of 2–3 particles (Figure 1C). An initial set of 1000 particles were manually localized to generate a preliminary cryo-ET map by subtomogram averaging (Methods). Iterative rounds of deep neural network-based particle picking^40,41^, followed by subtomogram averaging and classification, were used to localize particles across 355 tomograms of native cells (Figure S1A). In the final dataset, a major class consisting of 83% of the total 13,775 particles was resolved to 9.1 Å (Figures 1D, S1B-C and Table S1). The morphology of the adherent *M. pneumoniae* cells, with the larger membrane surface areas containing top and bottom views of the complex, resulted in an orientational bias of the particles and anisotropic resolution with a 3D Fourier shell correlation sphericity of 85% (Figure S1D-E)^42^.

**Figure 1.**
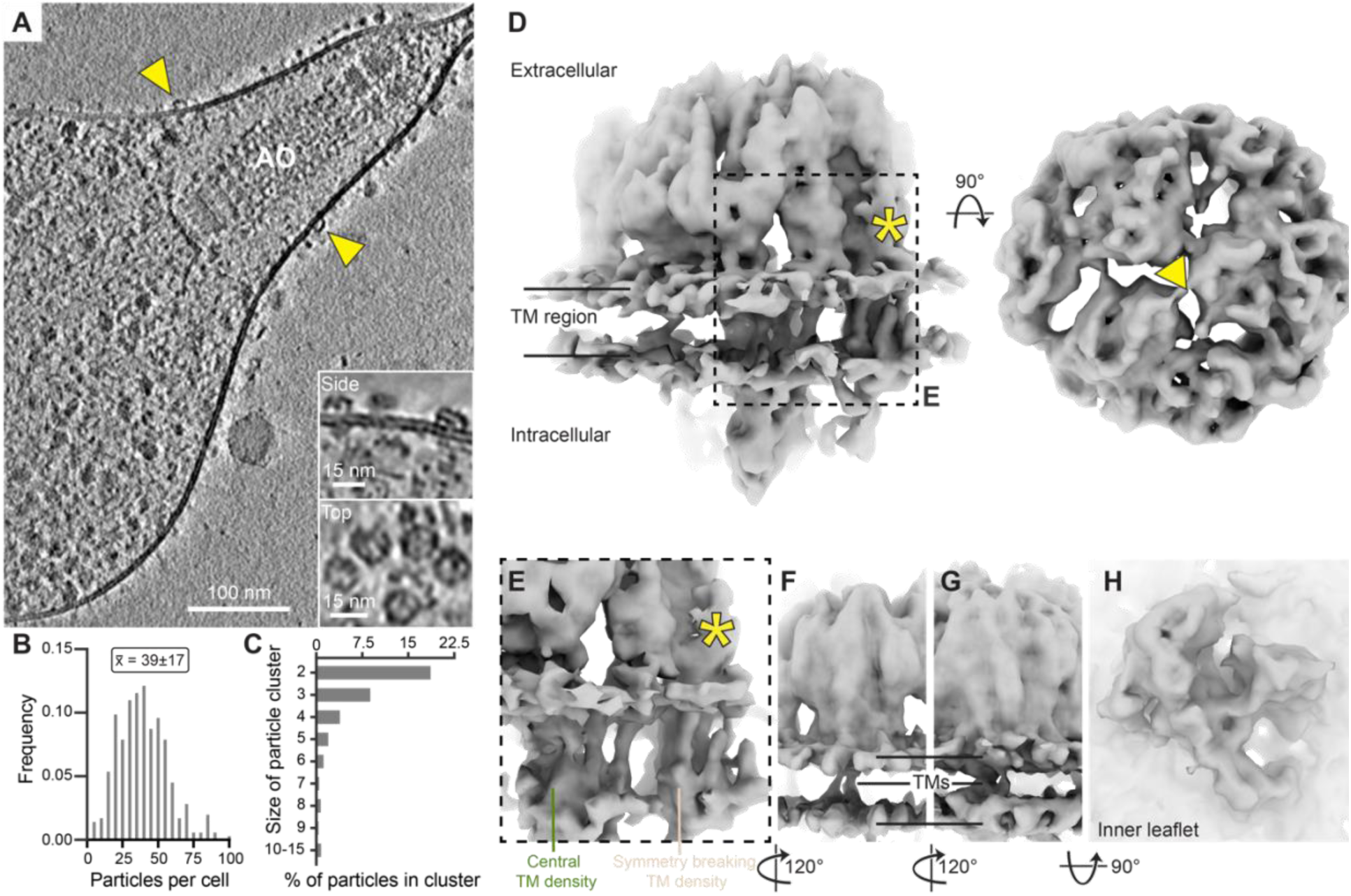
In-cell architecture of a cell-surface dome-like transmembrane complex. (A) Central tomographic slice of a *M. pneumoniae* cell. Yellow arrowheads indicate the extracellular dome-like complex. Insets show side- and top-views of the complex. The attachment organelle (AO) is indicated. (B) Histogram of particles per cell/tomogram (for 355 tomograms). Indicated are the mean ± standard deviation (SD). (C) Histogram of the percentage of particles observed in clusters of a given size. Particles are defined as being in the same cluster if the center-to-center distance is within 1.3 times the diameter of the particle. (D) Cryo-ET consensus map of the dome complex. The two lines indicate the membrane leaflets and the transmembrane (TM) region between them. The asterisk indicates density that breaks the three-fold symmetry and the triangle indicates the position of the pseudo-symmetry axis. (E) Enlarged view of the area indicated in (D), showing the two large transmembrane regions of the complex: the central (green) and symmetry-breaking (beige) densities. (F-G) The first (F) and second (G) minor transmembrane densities. (H) The cytosolic density of the complex. See also Figure S1.

The larger part of the density was extracellular and exhibited pseudo three-fold symmetry, giving rise to the characteristic dome-like appearance in the tomograms. The symmetry of the extracellular region was broken by an additional density present in only one of the asymmetric units (Figure 1D, *). This asymmetric density was associated with a transmembrane domain, located near an additional central transmembrane density within the dome (Figure 1E). Smaller densities spanned the plasma membrane at positions equivalent to the asymmetric extracellular density (Figure 1F-G). The cytosolic region consisted of a large continuous density with well-defined sub-domains (Figure 1H), which we observed to form multiple contacts with several transmembrane regions. To our knowledge, no complex with a similar architecture has been previously described.

### The extracellular dome is composed of large uncharacterized lipoproteins

To identify the proteins that constitute the extracellular part of the dome, we performed surface-shaving mass spectrometry (MS). In this approach, native cells were treated with exogenous proteases to selectively degrade surface-exposed protein regions. Cells were then lysed, digested and the resulting peptides labeled with tandem mass tags^43^ and analyzed by MS-based proteomics to compare their abundance in protease-treated and untreated cells, with the expectation that cell surface exposed proteins would have depleted peptides in the protease-treated conditions (Figure 2A, Table S2, Methods). We identified 113 putative surface proteins with significantly depleted peptides, including 55 proteins with predicted signal peptides and 32 proteins with predicted transmembrane regions^44,45^ (Figure 2A and Table S2). Only 43 of the identified proteins had an assigned or putatively-assigned function. To determine which of the candidates formed the dome, we extracted the extracellular region of the cryo-ET map using a cylindrical mask, predicted the structures of the determined surface-exposed proteins with AlphaFold2^46^, and rigid body fitted^47^ each of the predicted structures to the extracellular region of the map (Table S3, Methods). Among the candidates, three previously uncharacterized and homologous proteins MPN436, MPN444, and MPN489 (136-146 kDa), showed markedly higher cross-correlation values between the map and the models compared to other similarly sized proteins (Figure 2B and Table S3). This suggested that one, or all three proteins, make up part of the extracellular dome complex. Henceforth, we refer to these proteins as the major dome-forming proteins (Mdp) 436, 444 and 489. The Mdps are large lipoproteins with similar predicted structures, and each fitted equally well into either of the three asymmetric subunits of the density (Figure 2C and S2A-C). The predicted structures contained large disordered regions not resolved in the map, and many domain insertions, but could be divided into an N-terminal (N-ter) and C-terminal (C-ter) region (Figure 2C and S2D). While the sequence identity of the Mdps is only 19-23% (Figure S2E), their predicted structures share high similarity with a RMSD of 2.6-4.4 Å (over 690-855 Cα atoms), and close resemblance to the single particle cryo-EM structure of Mdp444^48^. In *M. pneumoniae,* all three proteins are essential under normal growth conditions, as determined based on a transposon-mutation screen^19^. They belong to the MG307/MG309/MG338 family of proteins, which includes eight members in *M. pneumoniae*. However, the remaining five proteins are truncated (Figure S2F), expressed at low levels^18^ and non-essential for growth^19^, making them unlikely candidates for the complex.

**Figure 2.**
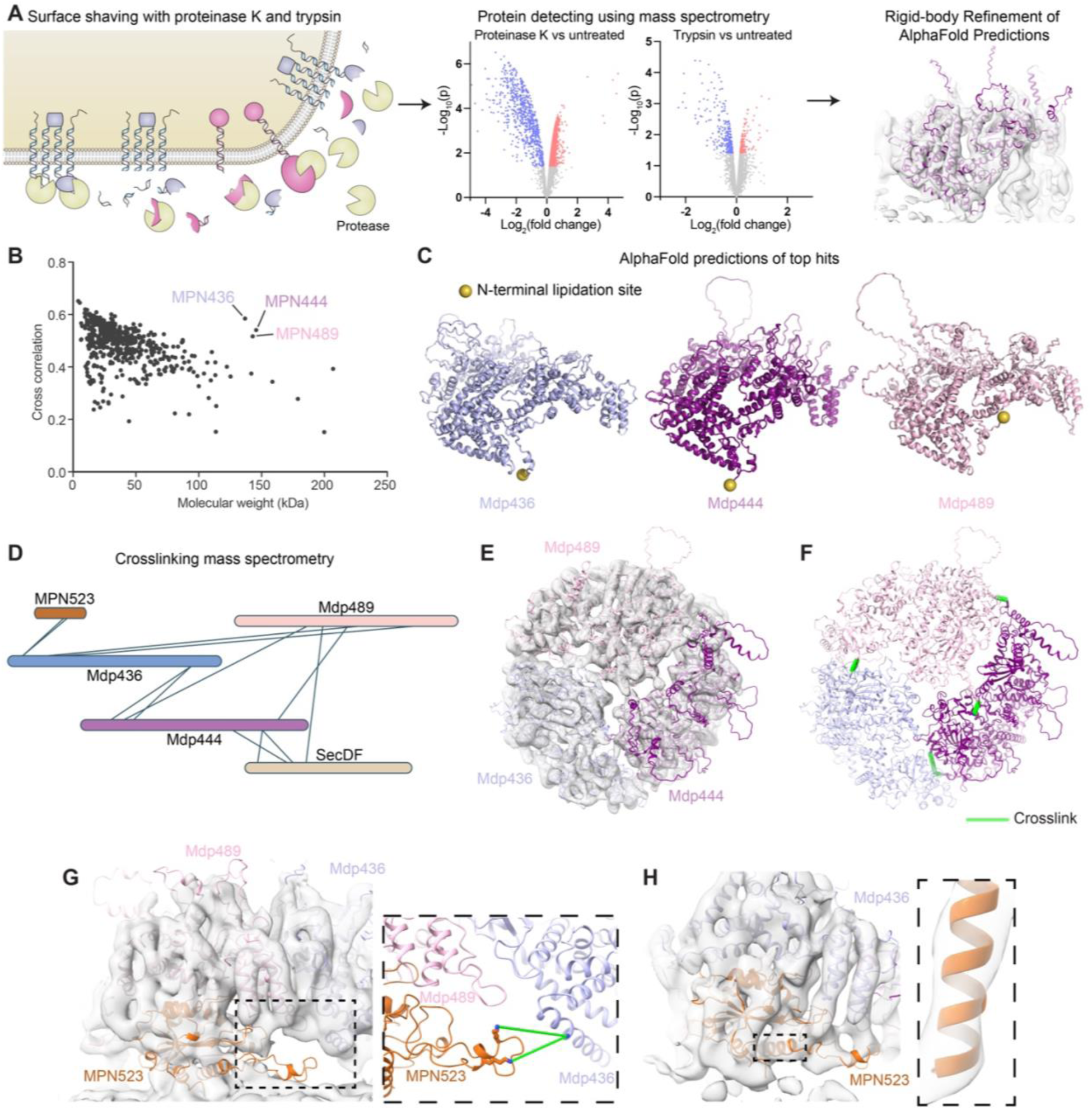
Identification of the extracellular lipoproteins forming the extracellular dome. (A) Workflow for identifying the extracellular proteins of the dome complex: cells were treated with the proteases proteinase K or trypsin, changes in abundance +/- protease treatment were used to detect shaved (depleted) proteins using mass spectrometry, and the AlphaFold2 predicted structures of the candidate proteins were systematically fitted into the extracellular regions of the cryo-ET map. (B) Plot of the cross-correlation for the systematic fitting between the rigid-body fitted models and the cryo-ET map, shown as a function of molecular weight. MPN436, MPN444, and MPN489 had substantially higher scores compared to other proteins of similar molecular weights. (C) The AlphaFold2 predictions of the top hits: Mdp436, Mdp444, and Mdp489 (previously MPN). The position of the N-terminal lipidation site is indicated with a yellow sphere. (D) In-cell crosslinking MS map of Mdp436, Mdp444, and Mdp489, MPN523, and SecDF. (E) Structural model of the complex between Mdp436, Mdp444, and Mdp489 fitted to the map. (F) The crosslinks between the Mdps are indicated by green lines, shown on the model. (G) Structural model of the complex between the Mdps and MPN523 fitted to the map. The inset shows the two crosslinks between MPN523 and Mdp436 as green lines. (H) The predicted multimer model of the Mdp436-MPN523 complex fitted to the map, suggests a second copy of MPN523 likely occupies this area. The inset shows the fit of the MPN523 residue 176-193 helix to the map at a higher map threshold. See also Figure S2.

The resolution of the cryo-ET map did not allow distinguishing between a homo- and hetero-trimeric arrangement of the Mdps. To resolve this, we reanalyzed a previously published in-cell crosslinking MS (CL-MS) dataset from *M. pneumoniae*^17^ using a different error cutoff (Methods). We identified crosslinks between all three Mdps, suggesting that they can form a heterotrimeric complex (Figure 2D). All self-crosslinks were consistent with the AlphaFold predictions (Figure S2A-C), and the intramolecular crosslinks suggested that a homo-trimer was unlikely. Moreover, mapping the crosslinks onto the protein models fitted into the map revealed a single possible arrangement that fully satisfies all inter-protein interactions (Figure 2D-F). Together, these findings strongly support a model where the C-ter of Mdp436 contacts the N-ter of Mdp444, the C-ter of Mdp444 contacts the N-ter of Mdp489, and the C-ter of Mdp489 contacts the N-ter of Mdp436 (Figure 2D-F).

In addition, we identified two crosslinks between Mdp436 and a fourth uncharacterized protein, MPN523 (Figure 2D). MPN523 is a 34 kDa essential lipoprotein, predicted to consist of a globular domain and a large disordered loop (Figure S2G; detailed bioinformatics in Figure S3). The location of the crosslinks indicated that MPN523 contacts the N-ter of Mdp436, and therefore likely has a larger interaction surface with the C-ter of Mdp489. We used AlphaFold2 to predict a multimer model of Mdp489, and MPN523 (Methods). The multimer prediction fitted well to the density corresponding to Mdp489 and an unassigned density the size of MPN523 (Figure 2G). The loop in MPN523, containing the crosslinked residues, extended towards Mdp436, supporting this spatial arrangement (Figures 2G, inset and S2H). This suggested that MPN523 is also part of the complex, positioned below the C-ter of Mdp489 and contacting the N-ter of Mdp436. Due to the pseudo-symmetry of the complex, a corresponding density was observed near the C-ter of Mdp436. The AlphaFold2 multimer prediction of the complex between Mdp436 and MPN523 fitted this region well, exhibiting well-resolved secondary structure elements (Figure 2H). This was in line with the abundance of MPN523 being 1.7 times higher than the Mdps measured in *M. pneumoniae* cells grown in minimal media^18^. In summary, we were able to putatively assign four uncharacterized proteins, consisting of a heterotrimer of the large Mdp lipoproteins, likely interacting with two copies of MPN523. The placement of these five proteins explained all extracellular regions of the complex.

### The major dome-forming proteins are distant homologs of PrsA

While the gene identity of the major complex constituents was now putatively assigned, the functional relevance of the complex remained unclear. The Mdps lacked functional annotation, with the only annotated domain (Interpro: IPR022186) being of unknown function and found almost exclusively in the *Mycoplasma* genus. Sequence-based homology searches were unsuccessful, likely due to the high mutation rate in *Mycoplasmas*^49^ and the presence of large disordered regions in the proteins (Figure 2C) which introduced gaps in the sequence alignments. Additionally, structure-based homology searches on the predicted models of the full-length Mdps, did not reveal any homologs of known function. We therefore used the predicted structures to define domains in the Mdps by removing disordered regions and insertions (Fig. 3A, Methods), and investigated each domain individually using the structure-based homology search algorithms DALI and FoldSeek^50,51^. This revealed that the N-ter helical domain of the Mdps exhibits homology to the trigger factor/SurA substrate-binding domain (Interpro: IPR027304) found in many chaperones, including trigger factor, SurA, and PrsA (Figure 3B-C and Tables S4-5). The homology search also showed that the domain starting between residues 222-229 in the Mdps was homologous to the peptidyl-prolyl cis-trans isomerase (PPIC) domain (Interpro: IPR000297, Figure 3B, D and Tables S4-5). This domain is often found in periplasmic or extracellular chaperones such as SurA and PrsA, and catalyzes the isomerization of proline between the trans and cis conformation, which is often the rate-limiting step in protein folding^52^. This domain organization is similar to that observed in the extracellular chaperone PrsA^53^, and is repeated in the C-ter (Figure 3B and Tables S4-5), suggesting that the Mdps evolved through an internal duplication. The C-ter PPIC domain indeed provided high scores in both DALI and FoldSeek for most of the Mdps, but not the C-ter trigger factor/SurA domain (Tables S4-5). We therefore used FATCAT^54^, where pairwise alignment confirmed that the N-ter and C-ter trigger factor/SurA domains were significantly similar across all three Mdps (Figure S3A-B, Methods). Combined, this analysis showed that all three Mdps contain an internal duplication, with each exhibiting a domain organization similar to that of the chaperone PrsA.

**Figure 3.**
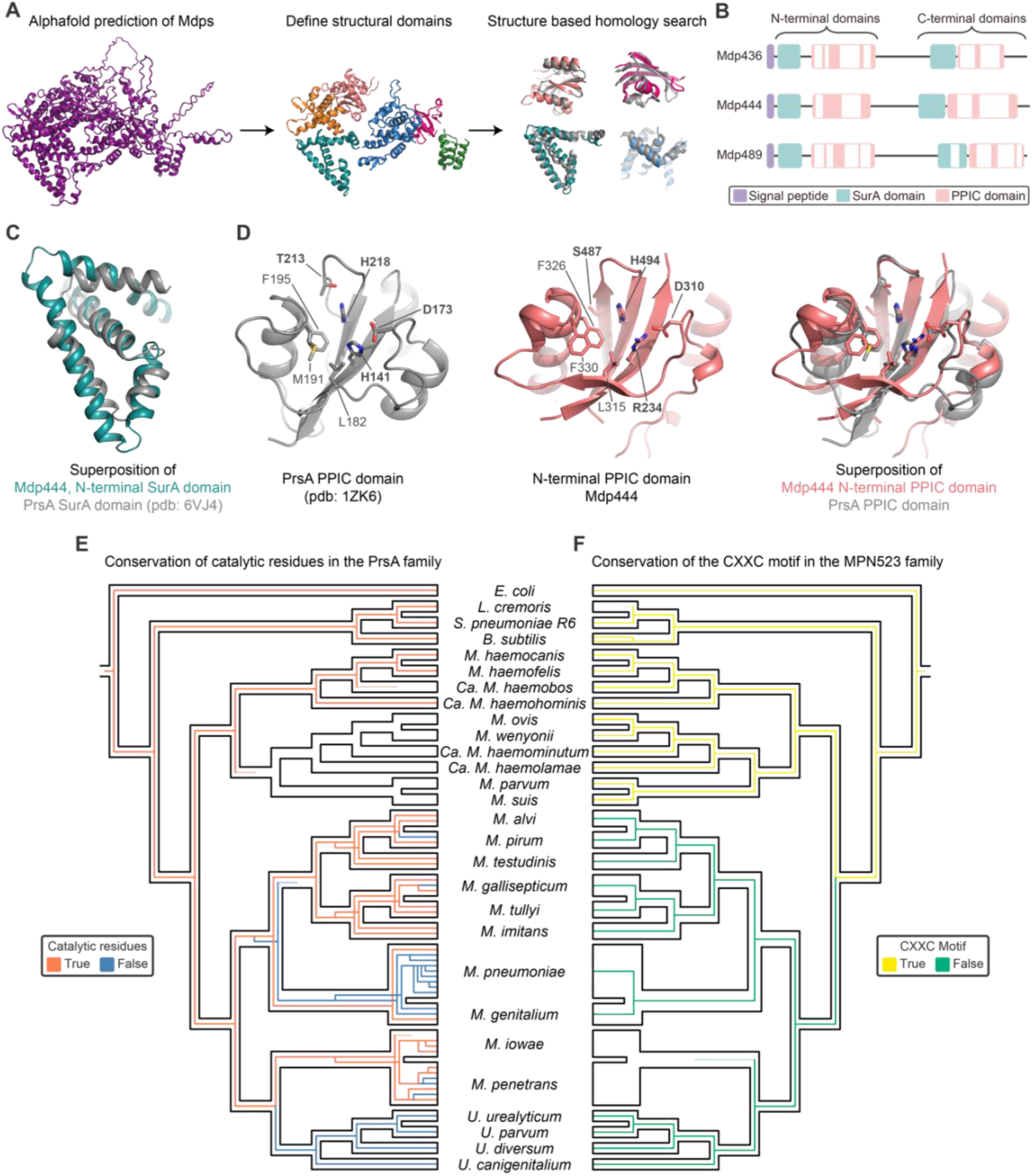
The major dome-forming proteins share remote homology with chaperones. (A) Workflow for determining homology of Mdps: structural domains of the Mdps were determined and excised from the AlphaFold2 predictions. These domains were used to perform structure-based homology searches using DALI and FoldSeek. Shown is the Mdp444 N-ter SurA domain (teal) aligned to pdb: 6VJ4 (grey), the N-ter PPIC domain (salmon) aligned to afdb: AF-A0A3M6KDR7-F1-v4 (grey), the C-ter SurA domain (blue) aligned to pdb: 5HTF (grey), and the C-ter PPIC domain (pink) aligned to afdb: AF-A0A5C6BMD5 (grey). (B) The domain arrangement determined from the structural homology searches shows that all three Mdps have an N- and C-terminal structural unit both consisting of a SurA domain followed by a PPIC domain. The white boxes indicate insertions. (C) Superposition of SurA domain from the Mdp444 N-terminal SurA domain (teal) and PrsA from *Bacillus anthracis* (pdb 6VJ4; grey). (D) The PPIC domain from *Bacillus subtilis* PrsA (pdb 1ZK6; grey) with the residues important for catalytic function indicated in bold, and the residues important for substrate binding indicated in regular text. The corresponding residues are indicated on the N-terminal PPIC domain of Mdp444, followed by a superposition of the two, showing that the catalytic and substrate binding residues are conserved. (E-F) A reconciliated phylogenetic tree, showing the evolutionary events of (E) the Mdp family of proteins, and (F) MPN523 in *Mycoplasma*, and other bacteria. The black outline indicates the species tree, and the colored lines indicate the proteins. The lines are colored to indicate whether (E) the catalytic residues are conserved (orange) or not (blue) for the Mdps, or (F) whether the CXXC motif is conserved (yellow) or not (green). The sequence similarity and alignment length of all *M. pneumoniae* paralogs of the Mdps is shown in Figure S2E-F. See also Figure S3.

However, the presence of the PPIC domain fold alone does not define enzymatic activity. We therefore investigated whether the catalytic and substrate-binding residues are conserved in the Mdps. For Mdp436 and Mdp489, the catalytic residues were not conserved (Figures S3C). However, structural alignment of the N-ter PPIC domain in Mdp444 and the PrsA PPIC domain from *Bacillus subtilis* revealed that most of the catalytic and substrate binding residues were conserved, with the exception of His141 (PrsA) which is substituted by Arg234 in Mdp444 (Figure 3D). The observation that only Mdp444 out of the three Mdps retains the catalytic residues, taken together with the almost equal expression level of the Mdps^18^, further supports a heterotrimeric arrangement of the complex. Next, we compared the amino acid positions corresponding to the catalytic residues in all proteins homologous to the Mdps in the *Mycoplasmatota* family using HMMER^55^ (Methods). This revealed that if a histidine and an aspartate were present in the positions equivalent to His494 and Asp310 in Mdp444, most proteins contained a conserved arginine in the position equivalent to Arg234 in Mdp444 (Figure S3C). Furthermore, this analysis showed that all species, except for the *Ureaplasmas,* possess at least one copy of the Mdp homolog that retains the catalytic residues (Figures 3E and S3C).

We used the same structure-based search strategy to find homologs of MPN523, which showed that it adopts the Thioredoxin-like superfamily fold (Interpro: IPR036249, Tables S4-5, Methods). The closest homolog identified was a thioredoxin-like protein containing the characteristic CXXC motif typically found in Dsb chaperones^56^ (Figure S4D). The DsbA family utilizes conserved cysteines to facilitate proper disulfide bond formation in their substrates. Interestingly, MPN523 has lost the two conserved cysteines and instead has a large inserted loop in the cysteine-containing helix (Figure S3E). Thus, while MPN523 may still retain the substrate-binding capacity of the thioredoxin-fold chaperones, it had lost the ability to catalyze disulfide bond formation. To investigate the evolutionary history of MPN523, we conducted a phylogenetic analysis (Figures 3F and S3F, Methods), which showed that YdfQ and BdbA in *B. subtilis* and DsbD in *E. coli* are orthologs of MPN523. The analysis also revealed that the CXXC-motif was lost after *Mycoplasma* speciation, separating the *Mycoplasmas* into two distinct groups (Figures 3F and S3F); most of the bacteria retaining the CXXC motif lost the Mdp genes, suggesting that the emergence of the dome complex and loss of CXXC may have co-evolved (Figure 3E-F).

In summary, our bioinformatical analysis revealed that the Mdps are homologous to the outer membrane chaperone PrsA, and that Mdp444 retained an enzymatically active PPIC domain. Furthermore, MPN523 shares homology to disulfide bond formation chaperones, but has lost the catalytic CXXC motif. Together, these findings suggest that the dome forms an extracellular folding chamber, which is likely involved in chaperoning membrane and secreted proteins.

### The Sec-translocon is an integral constituent of the dome complex

The CL-MS data^17^ indicated four crosslinks from the C-ter of Mdp444, and the N-ter of Mdp489, to the transmembrane protein SecDF (Figure 2D), suggesting that SecDF may correspond to the symmetry-breaking element of the complex. SecDF is a 108 kDa protein, consisting of 11 transmembrane helices and an extracellular domain, which can be divided into a head and base region (Figure S4A-B). The AlphaFold2 predicted models of the *M. pneumoniae* SecDF fitted each of the extracellular and transmembrane regions well, but the interdomain arrangement was incorrect. To resolve this, we separated the extracellular and transmembrane regions into two rigid bodies during refinement, which provided a good fit to the map and satisfied the four crosslinks to Mdp444 and Mdp489 (Figure 4A-D, Methods). The helix-extended loop-helix (HeH) motif near the Mdp444 C-ter interacted with the base of SecDF, while the first alpha-helical domain of the Mdp444 C-ter interacted with the SecDF head (Figure 4A-B).

**Figure 4.**
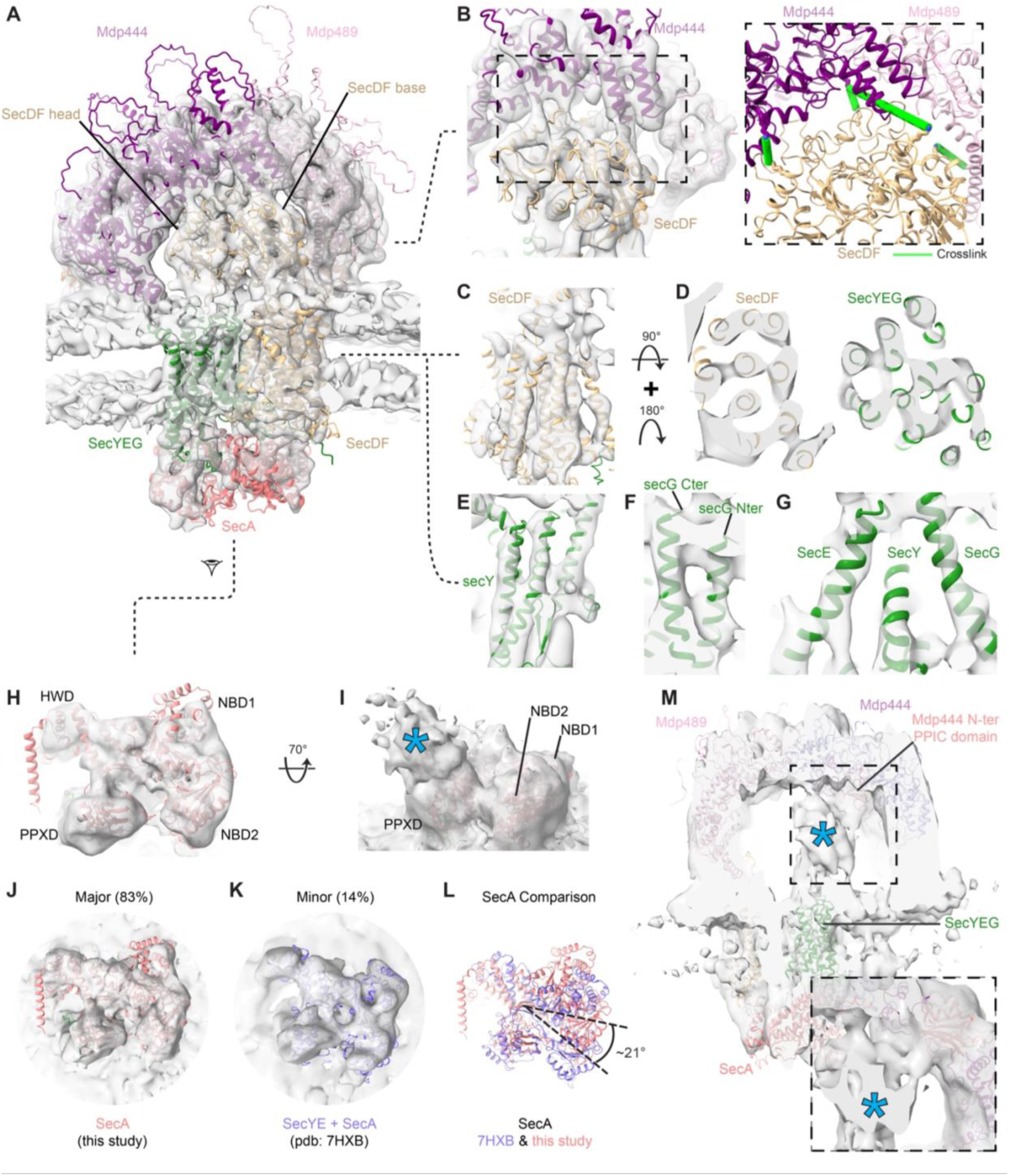
The Sec-translocon constitutes an integral part of dome complex. (A) The full structural model of the dome complex, including Sec proteins, the Mdps and MPN523, fitted to the cryo-ET consensus map. (B) Detailed view of the SecDF-Mdp444 interaction interface. The inset shows the crosslinks from Mdp444 and Mdp489 to SecDF as green lines on the structural model. (C-G) Detailed view of the transmembrane regions of SecDF (C-D), SecY (D-E), SecG (F), and SecE (G), fitted into the map. (H) Detailed view of the fit of SecA to the cytosolic density. The different domains of SecA are indicated. (I) SecA shown in the cryo-ET map contoured to a low threshold. Excess density on top of the PPXD is indicated by the blue asterisk, and likely corresponds to preprotein. (J-K) SecA in the major (J) and minor (K) conformations of the dome complex. (L) Structural alignment of the two SecA conformations, indicating a 21° rotation between the two. (M) The structural model of the dome complex fitted into the map contoured at low threshold. The N-terminal PPIC domain of Mdp444 is indicated in salmon, and the rest of Mdp444 is in purple. A putative substrate density (translocated protein) protrudes from the SecYEG channel into the dome (blue asterisk). The inset shows that the substrate interacts with the N-terminal PPIC domain of Mdp444. See also Figure S4.

SecDF is a cofactor in Sec-dependent translocation, which also includes the translocation channel SecYEG, and the cytosolic ATPase SecA^57^. To investigate whether the remaining densities could correspond to these Sec proteins, we predicted the structure of the predominantly transmembrane SecYEG and cytosolic SecA (92 kDa) (Figure S4C-D, Methods). The SecYEG model contained 10 transmembrane helices, which agreed well with the map (Figure 4D-G). The SecA model explained almost all of the cytosolic density of the map (Figure 4H). The density for the nucleotide-binding domain (NBD) 1 was less well resolved than the remaining SecA, and is known to undergo the largest domain movement during ATP hydrolysis^36^. When contouring the map at a lower threshold, we observed excess density not explained by our model near the preprotein crosslinking domain (PPXD) of SecA (Figure 4I, *). As the PPXD is responsible for preprotein binding^58^, this density may correspond to proteins being translocated. We observed that SecA adopted a closed conformation, as the PPXD was completely separated from the helical wing domain (HWD; Figures 4H and S4E). Fitting of SecA and SecY together showed that they have a large interaction surface formed mainly by the helical scaffold domain and PPXD, as previously described^37^ (Figure S4F). Notably, SecA was rotated 16-25° with respect to SecYEG as compared to previous *in vitro* structures^36,37,59,60^ (Figure S4F-H). Focused classification on the SecA density identified a minor SecA conformation resolved to 13 Å, which represented approximately 14% of the total particles (Figures 4J-K and S1B-C). Here, while the dome complex largely assumed the same conformation as the consensus map, SecA underwent a 21° rotation, resulting in a SecA-SecYEG arrangement similar to the *in vitro* structures (Figure 4L). Interestingly, we also observe a direct interaction between SecA and SecDF through the SecA HWD domain (Figures 4A and S4I).

On the extracellular side, we found SecDF and SecYEG to interact near the N-ter of Mdp444 (Figures 4A and S4K). The region comprising SecY α5, α6, and the loop connecting the helices (Ala215-Phe238), contacted a SecDF alpha helix (Ser472-Asn485) and nearby loop (Val209-Ala219). Although the resolution of the map prevented residue-level description of this interaction interface, several surface-exposed hydrophobic residues were modeled in these regions and are likely involved in the interaction (Figure S4J). Finally, we also observed a large heterogeneous density protruding from the exit tunnel of SecYEG into the dome cavity (Figure 4M, *), potentially corresponding to the translocated region of substrate protein. While the density is likely an average of different substrates at various stages of translocation, it was not completely randomly oriented within the dome, but interacted primarily with N-ter PPIC domain of Mdp444 (Figure 4M). This is in line with our bioinformatic analysis indicating that this domain of Mdp444 is catalytically active while those of the other Mdps are not (Figure 3D-E). Taken together, our analysis shows that the Sec-translocon, captured here in an actively translocating state, forms an integral part of the dome complex.

### The dome complex associates with the ribosome

Occasionally, we observed dome complexes in close proximity to ribosomes in the cellular tomograms (Figure 5A). To assess the significance of such co-localization, we analyzed the spatial organization of the dome complexes relative to all 70S ribosomes across 355 untreated cells (Methods)^10,17,20^, and found that 1.8% of dome complexes localized within 10 nm of the nascent peptide exit site of a ribosome (Figure 5B). Hypothesizing that the dome-ribosome association may depend on co-translational protein translocation, we extended our analysis to cells treated with antibiotics that stall protein synthesis (Figures 5B-E and S5), either during translation elongation using chloramphenicol (Cm), or transcription using pseudouridimycin (PUM)^10,17,20,61^. We indeed found that the fraction of dome-ribosome interactions markedly increased under these conditions to 8.5% in PUM, and 15.9% in Cm treated cells respectively (Figure 5B).

**Figure 5.**
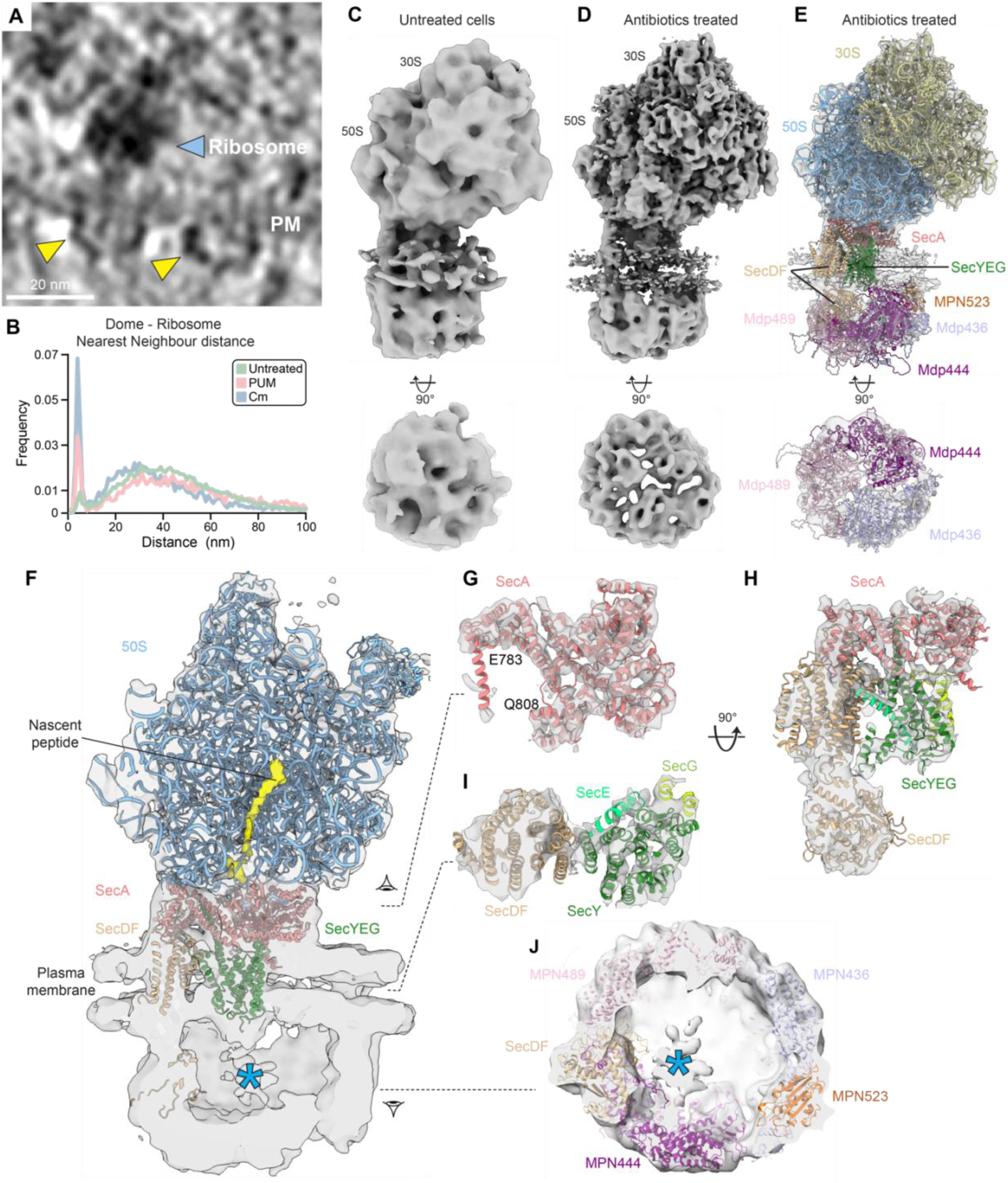
Functional association of the dome complex with the ribosome. (A) Tomographic slice of a *M. pneumoniae* cell showing a ribosome (blue arrowhead), the plasma membrane (PM) and the dome complexes (yellow arrowheads) on the cell surface. (B) Nearest neighbor analysis, showing the distance for each dome complex to the nearest ribosome in untreated cells and in cells treated with pseudouridimycin (PUM) or chloramphenicol (Cm). The distance was measured from SecA to the peptide exit site. (C-E) Cryo-ET maps of the dome-ribosome supercomplex resolved in untreated cells (C), or antibiotics-treated cells (D) and the corresponding structural model (E). (F) View of the nascent peptide density extending from the ribosome peptidyl transfer center to the exit site (yellow). Low density thresholding was used to visualize the substrate density inside the dome cavity (blue asterisk). (G-J) Model fitting of SecA (G), SecDF-SecYEG (H-I) and the Mdps (J) in the multibody derived maps. See also Figures S5 and S6.

To structurally characterize this interaction, we extracted ribosome subtomograms using a box size optimized to capture surrounding complexes (Methods). Ribosomes were first sorted based on the proximity to a nearby dome, followed by focused classification on the peptide exit region (Figure S5A). In untreated cells, we found 0.19% of 70S ribosomes in ribosome-dome complexes (151 particles), yielding a 19 Å resolution map (Figures 5C and S5B-C, Methods). In antibiotic treated cells, similar analysis identified 3% of 70S ribosomes to be in ribosome-dome complexes (2304 particles). Overall, the maps from untreated and treated cells exhibited the same architecture, except for different rotations of the 30S small ribosomal subunit due to the effect of the specific antibiotics (Figure S5B-G).

The increased occurrence of ribosome-associated dome complexes in cells treated with protein synthesis inhibitors indicated that translocation becomes stalled by the unterminated polypeptide chain emerging from the ribosome, which likely pulls the ribosome closer to the dome, eventually leading to stabilization of an otherwise transient supercomplex. To better resolve its structure, we pooled 2,304 supercomplexes from cells treated with different antibiotics^10,17,20^ (Figure S5G-J) and performed multi-body refinement. This resulted in a map with the dome complex resolved to 11 Å and the 50S large ribosomal subunit together with SecA resolved to 8.2 Å (Figures 5D and S5H-J).

We next rigid-body fitted the models of the *M. pneumoniae* ribosome^20^, the dome and the Sec-translocon into this map (Figure 5E-J, Methods). The ribosome and the SecA density could be unambiguously modeled, down to the secondary structure level details (Figure 5G), showing that their interaction is mediated by ribosomal proteins L23, L24 and L29 surrounding the peptide exit site (Figure S6A-C). Our model markedly differed from the ribosome-SecA complex reported *in vitro*^63^ in the orientation of SecA and its interaction interface with the ribosome (Figure S6D). The SecY-SecA orientation was similar to the minor SecA open conformation observed in the untreated cells, indicating a functionally active state (Figures 3J-L, S3F-H and S6E-G). SecDF and SecYEG did not appear to form any direct contacts to the ribosome, and no further unexplained densities corresponding to trigger factor or other factors was observed (Figure 5F, H). Except for SecA, the remaining dome complex conformation was similar to that in the free dome complex. Of note, a density for the nascent peptide could be mapped from the ribosome peptidyl transfer center to SecA (Figures 5F and S6A). We also observed a substrate density protruding from the SecYEG channel into the dome cavity (Figure 5F, J).

These maps provide the first direct structural evidence of SecA-mediated co-translational translocation. They further indicate that both post- and co-translational translocation can occur through the dome complex.

### SecDF undergoes structural rearrangement upon substrate and SecA release

Our results so far suggest that the dome acts as an extracellular folding chamber for newly translocated proteins. However, it remained unclear how a folded substrate can be released. We first examined the structural state of the dome complex in the antibiotic-treated cells where protein synthesis is stalled. Classification revealed that while most complexes retain a similar conformation to that observed in the untreated cells, a minor class resolved to 17 Å exhibited an opening of the dome (Figures 6A-D and S7A, Methods). In the Cm- and PUM-treated cells, this class corresponded to 28 % and 11 % of the complexes, respectively, and we did not observe any instances of this conformation in complex with the ribosome. We were also able to identify this open conformation in the untreated cells by performing a series of classifications first focused on SecDF, followed by SecA, and found it to only account for 1.2% of the complexes (Figure S1B-C). This analysis resolved a number of additional states (discussed below). In the open conformation, the extracellular domains of SecDF were rotated towards the center of the dome, and both the interactions of Mdp444 with the base and head regions of SecDF were lost. This resulted in an opening between Mdp444 and Mdp489 (Figure 6A-C), correlated with loss of density for the cytosolic ATPase SecA, and the substrate inside the dome cavity (Figure 6A, C-E). This suggested that by inhibiting protein synthesis, we enriched for and resolved a resting state of the complex, where no active translocation occurs.

**Figure 6.**
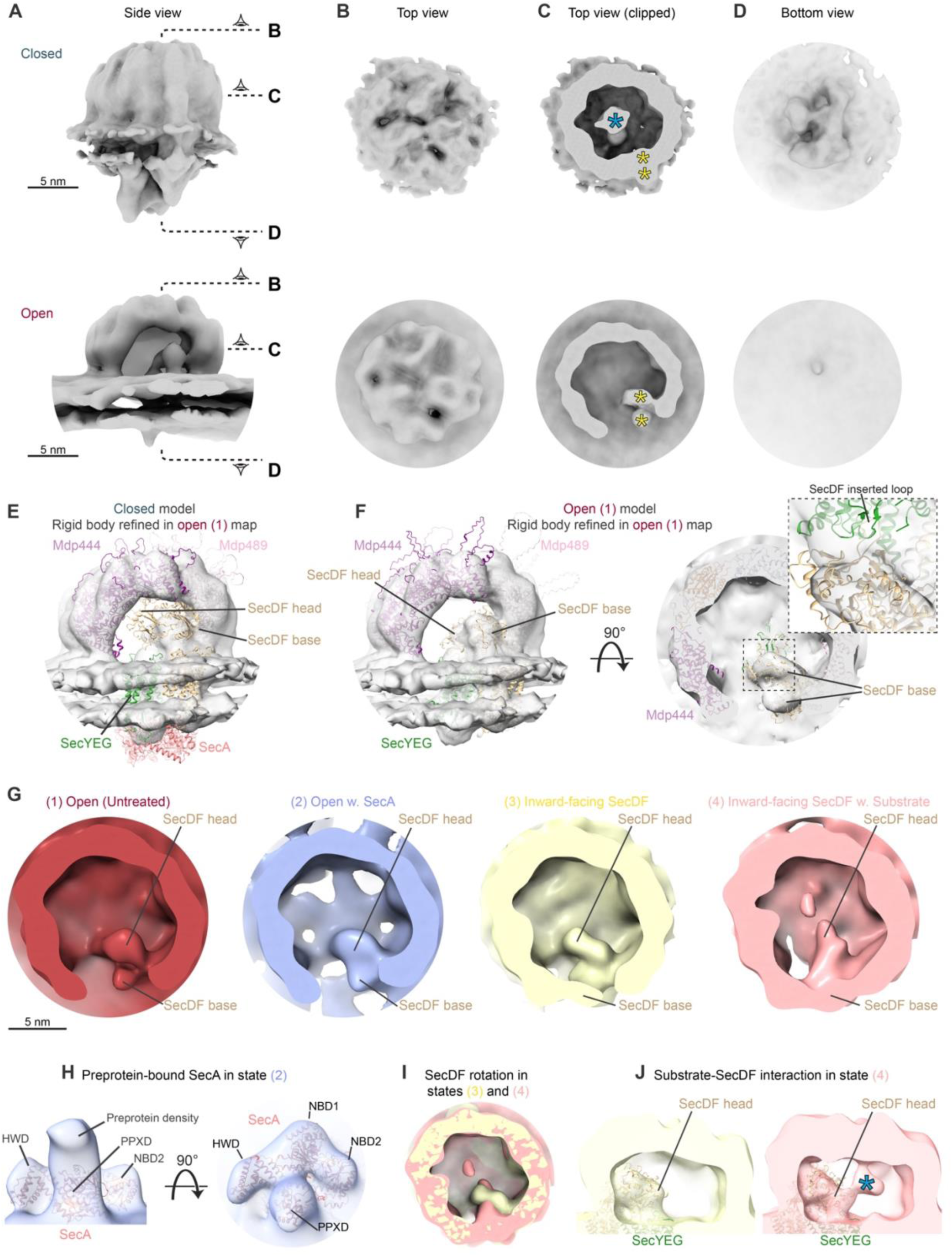
Structural rearrangement of the dome complex during translocation and substrate release. (A-D) Cryo-ET map of the closed dome complex conformation, low-pass filtered to 20 Å, for comparison with the map of the open conformation, shown as (A) side views, (B) top views, (C) clipped top views, and (D) bottom views. In the open conformation, an opening in the dome is observed, the substrate (blue asterisk) and cytosolic densities are absent. The SecDF head and base domains are indicated by yellow asterisks. (E) Map of the open conformation fitted with the structural model of the closed conformation. (F) Map of the open conformation after rigid-body refinement of models. The inset shows a detailed view of the SecDF head, and the loop inserted into SecYEG. (G) Views of the inner dome region in maps of the four states classified in untreated cells. Densities corresponding to the extracellular regions of SecDF are indicated. (H) The cytosolic density of the map of the open confirmation with SecA fitted shows density for the preprotein above the PPXD domain. (I) Cryo-ET maps of the inward-facing SecDF without (yellow) and with (pink) substrate. SecDF undergoes a small conformational change. (J) The map of the inward-facing SecDF without and with substrate (blue asterisk, *) shown at a low threshold, with SecDF and SecYEG fitted, showing that SecDF contacts the substrate density in the substrate containing map. See also Figure S7.

To better understand which proteins undergo conformational changes between the open and closed states, we fitted the model of the actively translocating conformation into the map of the open conformation (Figure 6E). This confirmed that SecA was absent in the open state. In addition, SecDF, SecYEG and Mdp444 underwent conformational changes (Figure 6E-F and Figure S7B-D). Interestingly, a predicted AlphaFold3^64^ model of the SecYEG-SecDF complex fitted the open conformation, but not the closed one (Figures 6F and S7B). We therefore rigid-body refined the subcomplex into the map of the open conformation, and performed refinement of Mdp444 separately as a single rigid body (Figure 6F and Methods). Our structural model showed that transitioning from the closed to open conformation, the extracellular domain of SecDF undergoes a 28° inward rotation and 19 Å shift. This movement created an opening in the extracellular dome, presumably providing translocated and folded substrates with an exit site (Figures 6A, E-F and S7E). The head of SecDF was now positioned above SecYEG (Figures 6F and S6B) and the loop of SecDF (Val209-Ala219), which interacted with SecYEG α5 and α6 in the closed conformation, had inserted into the exit tunnel of SecYEG (Figures 6F and S7B). This loop is likely responsible for accepting the incoming substrate during early translocation and initiates the proposed pulling action of SecDF^31,65^. Alongside the large conformational changes observed in the extracellular domain of SecDF, the transmembrane domains of SecYEG and SecDF had moved closer together (Figure S7B-D). Furthermore, Mdp444 tilted slightly away from the membrane, increasing the opening between the C-ter of Mdp444 and N-ter of Mdp489 from approximately 5 Å to 10 Å (Figure S7E, Methods). Further classification of the particles from the open conformation revealed substantial variability in the size of the opening between Mdp444 and Mdp489, extending to approximately 24 Å in the widest conformation (Figure S7E). This flexibility may be necessary to accommodate the release of larger substrates.

As mentioned above, we observed several additional states in the untreated cells that were distinct from the open (1) conformation (Figures 6G and S1B). Although all three maps were resolved to modest resolutions (25-29 Å, Figure S1B-C), they showed conformational differences in SecA, SecDF and Mdp444. The second state (2) closely resembled the open state, with the key difference being the presence of SecA in state (2) (Figures 6G and S7F). We did not observe any substrate density inside the dome, but an additional density above the PPXD of SecA likely corresponds to bound preprotein (Figure 6H). Thus, this conformation may represent a pre-rather than a post-translocation state of the complex. Comparison of the position of SecA to the closed conformation showed that SecA adopted the same overall conformation, but was rotated to accommodate the changed positions of SecYEG and SecDF in the open state (Figure S7G). In a third state (3), we similarly observe SecA bound to a putative preprotein density and SecDF in an inward-facing conformation as in the open state (Figures 6G and S7H-J). Here, the interaction between the base of SecDF and Mdp444 was intact (Figures 6G and S7H), and there was a change in the orientation of the head of SecDF (Figure S7I). No substrate density was found inside the dome (Figures 6G, I and S7I). In a fourth state (4), Mdp444 interacted with the base of SecDF, but the head region was still facing inward, in close resemblance of state 3 (Figures 6G and S7K). In contrast to state 3, we now resolved a substrate density inside the dome, which was continuous with the SecDF head density (Figure 6I-J). The SecDF head was rotated, likely due to its interaction with the substrate (Figure 6I). The substrate density was nevertheless weaker than in the closed conformation (Figure 6C, G, I-J), suggesting an early state of translocation.

Taken together, these states allowed us to propose a model for the translocation of unfolded proteins across the membrane in *Mycoplasma* (Figure 7A). Initially, in its open resting state, the SecA-unbound dome complex positions the SecDF head above the exit tunnel of the SecYEG channel. The recruitment of preprotein-bound SecA to the complex facilitates the interaction of the SecDF base with Mdp444. Protein translocation initiates through the SecYEG channel and the N-terminus of the translocated protein is recognized by SecDF, in a manner likely driven by the proton-motive force. Next, the SecDF head interacts with Mdp444, likely facilitating substrate handover. This leads to the closed conformation of the complex, enabling SecA-driven translocation through ATP hydrolysis. The translocated substrate is folded within the dome lumen, aided by the extracellular chaperones of the dome, and SecA dissociates from the complex. Finally, the SecDF-Mdp444 interaction is lost, leading to the opening of the dome and substrate release.

**Figure 7.**
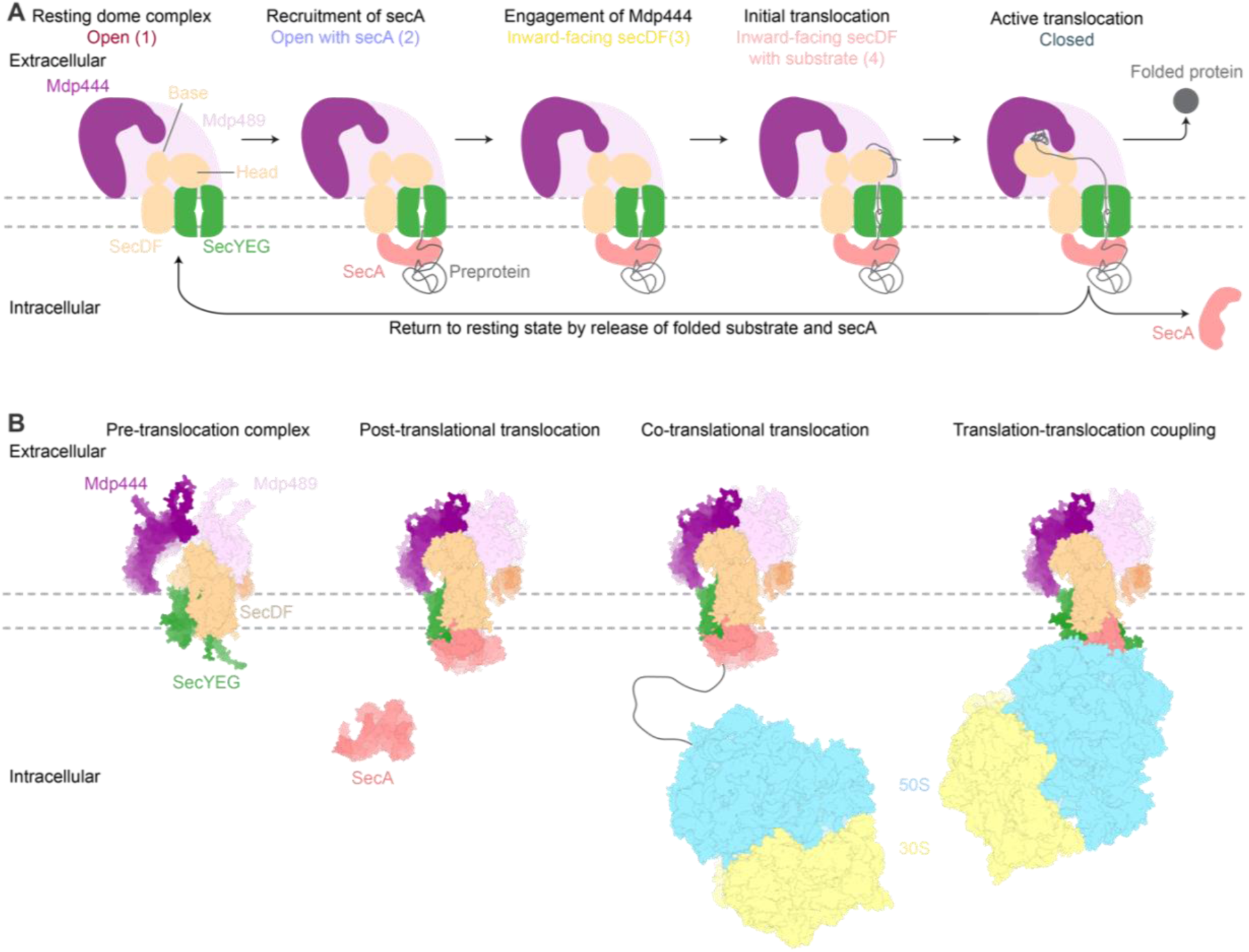
Models of protein translocation and folding by the non-canonical dome-translocon complex. (A) Schematic model of the translocation cycle in *M. pneumoniae*. SecA binding primes the complex for translocation, involving reorganization of the SecDF-SecYEG architecture, and interactions with the extracellular dome proteins. (B) Structural models of the different dome-complexes. In the pre-translocation state, the SecA-unbound dome is in a resting state. Association of SecA with a substrate initiates translocaiton. Co-translational translocation can either happen in a flexible form with ribosome and dome associated via the nascent peptide or directly through physical coupling of the two macromolecular complexes, forming a supercomplex.

## Discussion

In this study, we demonstrated the feasibility of determining the identity and function of a previously uncharacterized membrane-bound protein complex by in-cell cryo-ET. This required the integration of multiple publicly available data resources, including CL-MS and quantitative proteomics, bioinformatics methods for protein structure prediction, sequence and structure alignments, alongside tailored analyses of the cryo-ET density, surface-shaving proteomics, and cellular perturbations. Similar multi-modal approaches have recently been applied to well-characterized systems, such as the flagella apparatus^12,14^. A key challenge here was inferring the function of the uncharacterized extracellular proteins, complicated by limited annotation of membrane proteins^22^ and the high mutation rate in *M. pneumoniae*^49^, which hampered sequence-based homology-searches. We therefore relied on structure-based homology searches for isolated protein domains, which enabled functional assignment of the previously uncharacterized proteins. Although our investigation began with visualization of an unknown extracellular complex, we discovered that the conserved Sec-translocation machinery formed the majority of the cytosolic and transmembrane components of the complex. This ultimately allowed us to resolve a sub-nanometer resolution map and construct a full structural model of the bacterial holotranslocon, detailing its architecture, and the interaction interfaces between the Sec proteins *in situ*.

### Sec-translocon architecture

Although X-ray crystallography-derived structures of the SecDF complex are available^31,38,39^, its interactions with the other Sec proteins have not been described. Our study provides insights into the conserved interactions between SecYEG and SecDF. We found these interactions to vary between the open and closed conformations of the extracellular dome, but the Val209-Ala219 loop of SecDF was important in both interaction interfaces. Interestingly, we also found a direct interaction between SecA and SecDF. This interaction may be critical for coordinating the transition between different states during initiation and termination of translocation. SecA is known to dimerize during preprotein binding and recruitment to the membrane^66^. However, the oligomeric state of SecA during active translocation has been controversial^66–75^. In *M. pneumoniae,* we only observed SecA monomers interacting with SecYEG both during initiation and active translocation. The SecA-SecYEG subcomplex assumed two major conformations in our data. In the ribosome-bound state and what we term the SecA minor conformation of the free dome complex, the SecA-SecYEG subcomplex architecture was similar to the *in vitro* determined structures mimicking active translocation^36,37^. The major conformation found in the free dome complex deviated substantially from all previously determined structures^36,37,59,60^. Whether this alternate conformation reflects active translocation or is related to the association of other components of the complex, such as the extracellular chaperones or SecDF, will require further investigation.

The cellular copy number of SecY closely matches that of the Mdps in *M. pneumoniae*^18^, suggesting that the majority of SecY is bound within the dome complex. The Sec-translocon is the only protein translocation or secretion system identified in the *M. pneumoniae* genome^23,24^. We therefore hypothesize that most lipoproteins, secreted proteins, and integral membrane proteins are processed by the dome complex, consistent with all the dome proteins being essential. Although YidC is expected to associate with the Sec-translocon for insertion of small membrane proteins^76^, we were unable to detect a corresponding density despite extensive classification efforts. Whether this is due to the complex being rare, given that YidC is expressed at one-third the level of SecY^18^, or because the complex is transient, remains to be seen. YidC can also function independetly of the Sec proteins, and has been shown to be sufficient for insertion of several membrane proteins^77–81^. It is thus possible that the reductive evolution of *Mycoplasma* may have driven YidC to function primarily independently of Sec in *M. pneumoniae*.

### Protein folding during translocation

We showed that the Sec-translocon forms a complex with a putative extracellular folding machinery in a nearly obligate manner in *M. pneumoniae*. Previous studies have shown that the Sec-translocon can interact directly with chaperones such as PpiD and SurA in *E. coli*^82–84^, and that the YfgM/PpiD complex can mediate handover to periplasmic chaperones^85^. We found that the Mdps are homologous to the Gram-positive chaperone PrsA, usually involved in folding of secreted proteins and which constitutes a bottleneck for their overexpression^86,87^. Interestingly, partial folding of the substrate during translocation has been suggested to help avoid backsliding in the SecYEG channel between SecA strokes^88^. Close association between the Sec-translocon and extracellular chaperones may therefore enhance translocation efficiency. This raises the question of why *Mycoplasma* evolved a more tightly controlled environment for folding and maturation of secreted proteins, provided by the dome. One possible explanation could be the absence of the cell wall or periplasmic space in *Mycoplasmas*, where secreted proteins in other bacteria would typically be folded before subsequent export to the cell surface. The lack of a cell wall helps pathogenic *Mycoplasma* evade detection by the host’s innate immune system^89^, but at the expense of exposing the bacterial surface proteins to the adaptive immune system. Therefore, the protective environment of the dome may ensure nascent proteins are fully folded and functional before being directly exposed to host defense mechanisms. This system may represent a specific target for future therapeutics, due to it being uniquely present in *Mycoplasmas*.

### Functional flexibility

Cryo-ET of intact cells captures macromolecules in action, unless the system is halted. This means that most complexes exhibit functional dynamic, complicating structural analyses and limiting the achievable resolution. This rich information nevertheless allows probing the conformational landscapes of macromolecular complexes. Recent in-cell cryo-ET studies provide detailed insights into the action of large macromolecular machines, including ribosomes^20,90–95^, chaperones^96,97^, and ATP-synthase^98^. Here, we demonstrate the feasibility of analyzing systems that are less abundant, and that exhibit only small conformational changes along their functional cycle. It would have been impossible to understand how a substrate could be released after translocation and folding by the dome complex from the analysis of the major, well-resolved, closed conformation alone. Comprehensive focused classifications revealed a rare (transient) resting state where a large inward rotation of SecDF and tilting of Mdp444 leads to an opening in the extracellular dome. This opening potentially facilitates substrate release after protein folding and signal peptide cleavage. Interestingly, while *M. pneumoniae* does have signal peptidase I activity, we and others have been unable to identify the corresponding enzyme^24,99,100^. We also did not observe any density near the signal peptide site in our cryo-ET maps, suggesting that signal peptidase I is either small or only transiently associates with the complex. We additionally identified multiple, low abundance pre-translocation states of the complex that allowed us to suggest a model for translocation initiation (Figure 7A). Previously, the following model based on X-ray structures of SecDF was proposed^31,38,39,65^: in a resting state of SecDF, the head region is inclined towards SecYEG, ready to interact with the preprotein emerging from the channel. The model suggests that binding of the substrate to the SecDF head triggers a transition, wherein the SecDF head, still inclined towards SecYEG, undergoes an upward rotation with respect to the plane of the membrane. This is similar to the transition we observed from the open conformation (state 1) to the intermediary states (3 and 4), which show an inward-facing SecDF. SecDF is then proposed to transition to a final state where the head rotates further and opens up the proton gate. The energy provided by the proton motive force allows for the release of the substrate, and resetting of SecDF back to the resting state^65^. Since our maps were not resolved to residue-level, we cannot determine the states corresponding to proton channel opening and closing. While our findings align with the model proposed by Tsukazaki et al.^65^ on many points, clear differences exists in the overall conformations, and future in-cell studies across diverse organisms will likely determine whether these differences reflect mechanistic variations between bacteria.

### Coupling of protein translation, translocation and folding

Our findings suggest that the dome complex can participate in both post- and co-translational translocation (Figure 7B). Co-translational translocation may occur in the form of flexible tethering of the ribosome via the nascent peptide or by direct physical interaction between the two complexes. We observed a stable ribosome-dome complex at low abundance in untreated cells (1.8%), and an enrichment of this complex following cell treatment with protein synthesis inhibitors (8-16%), indicating that some degree of physical translation-translocation coupling must occur in *M. pneumoniae*. The maps of the stable complex show association of the dome with the ribosome through SecA and provide direct structural evidence of SecA-mediated co-translational translocation in cells. Furthermore, the nascent peptide density traced through the entirety of the peptide exit tunnel to SecA indicates direct handover from the ribosome to SecA (Figure 5F). However, the enrichment of the ribosome-dome complex during antibiotic treatment also suggests that a substantial fraction of translocation is not strictly functionally coupled to translation. This enrichment could be a result of unperturbed ongoing translocation while protein synthesis is halted, leading to the ribosome being pulled to the translocon. Our findings align with growing evidence from *E. coli* showing that SecA-mediated co-translational translocation can occur both with and without physical coupling^26,32,33,63,101^.

## Conclusion

We demonstrated how integrating experimental and computational approaches enabled in-cell discovery of a previously uncharacterized protein complex, while at the same time uncovering new mechanistic aspects of conserved molecular machinery. We presented the first detailed structural analysis of the bacterial holotranslocon and provided direct structural evidence of coupled, SecA-mediated, co-translational protein translocation. By visualizing a range of pre-translocation conformations of the complex, we also suggested a mechanism for translocation initiation in *Mycoplasmas*. The observed tight association between the extracellular chaperone and the Sec-translocation system may have arisen due to the parasitic lifestyle and reductive evolution of *M. pneumoniae*, but could also represent a universal mechanism to increase translocation efficiency by mitigating substrate backsliding through the channel. Whether similar mechanisms are employed in other organisms remains to be explored. In summary, our work establishes a framework for harnessing cryo-ET to characterize novel macromolecular complexes of unknown function through direct visualization *in situ*.

## Acknowledgments

We thank the EMBL Proteomics and Electron Microscopy Core Facilities, the EMBL cryo-EM platform, EMBL IT, and especially Thomas Hoffmann for computational support. We thank the Mahamid group members for discussions, especially Lucas Adam and Evgenia Zagoriy for technical assistance. We also thank Kausthubh Ramachandran for helpful discussion regarding phylogenetic analysis. This work was supported by EMBL core funds. R.K.J was supported by the Independent Research Fund Denmark (grant number 0164-00010A). L.X. acknowledges support from the National Key Research and Development Program of China (2024YFA1306200). J.R. acknowledges funding by the Deutsche Forschungsgemeinschaft (DFG, German Research Foundation) under Germany’s Excellence Strategy – EXC 2008 – 390540038 – UniSysCat and project no. 426290502. J.M. acknowledges the support from an EMBL Infection Biology Transversal Theme Synergy grant, and a Chan Zuckerberg Initiative grant for visual proteomics (2021-234620).

## Author contributions

Conceptualization, R.K.J., L.X., M.Z.K, P.B. and J.M., Investigation, R.K.J., L.X., F.M., J.C.S., J.S., and S.L., Methodology, J.S., A.T., J.R., M.S., J.K., M.Z.K, P.B., J.M., Writing - Original Draft, R.K.J., L.X. and J.M., Writing – Review & Editing, All authors.

## STAR*Methods

### Key resources table

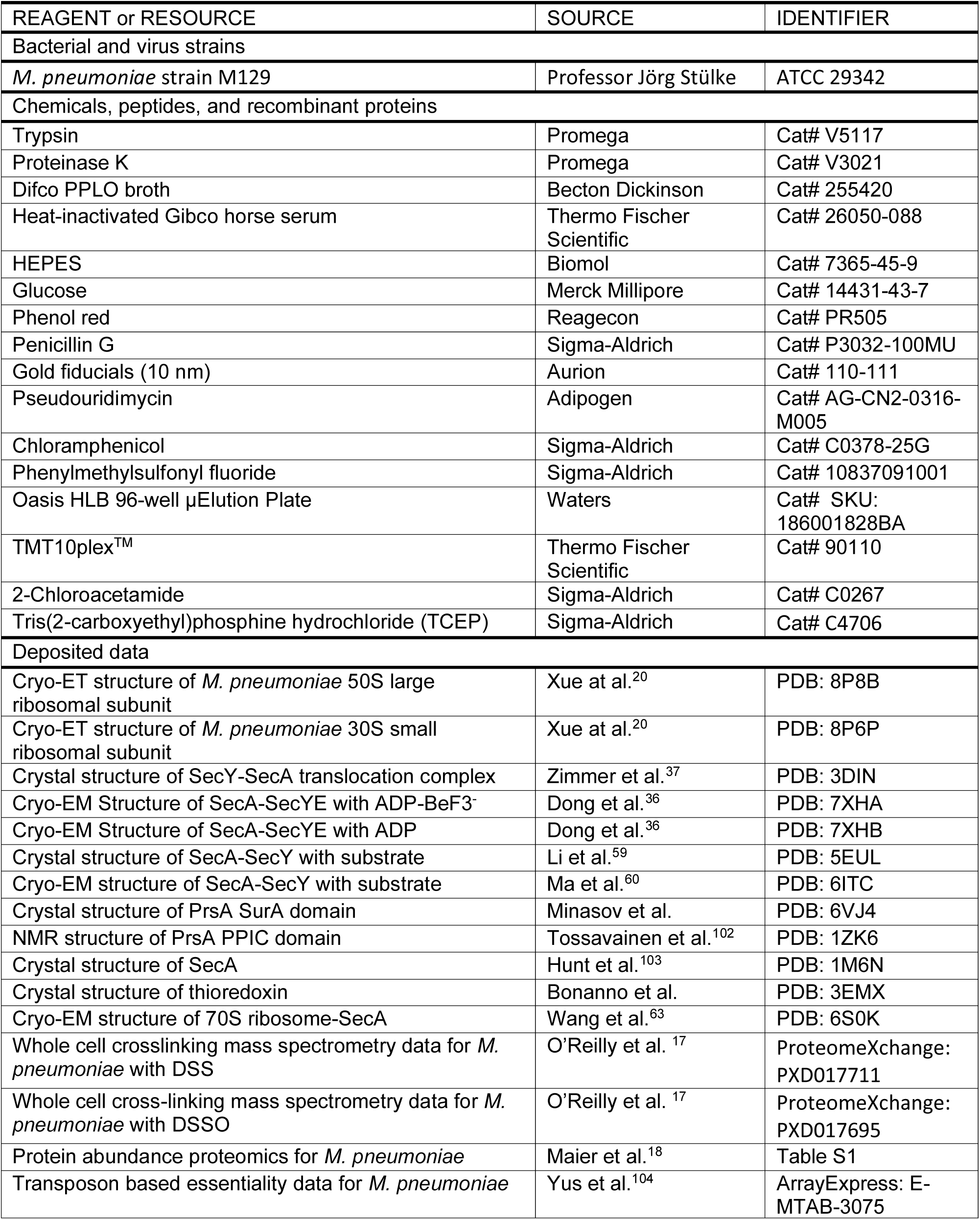
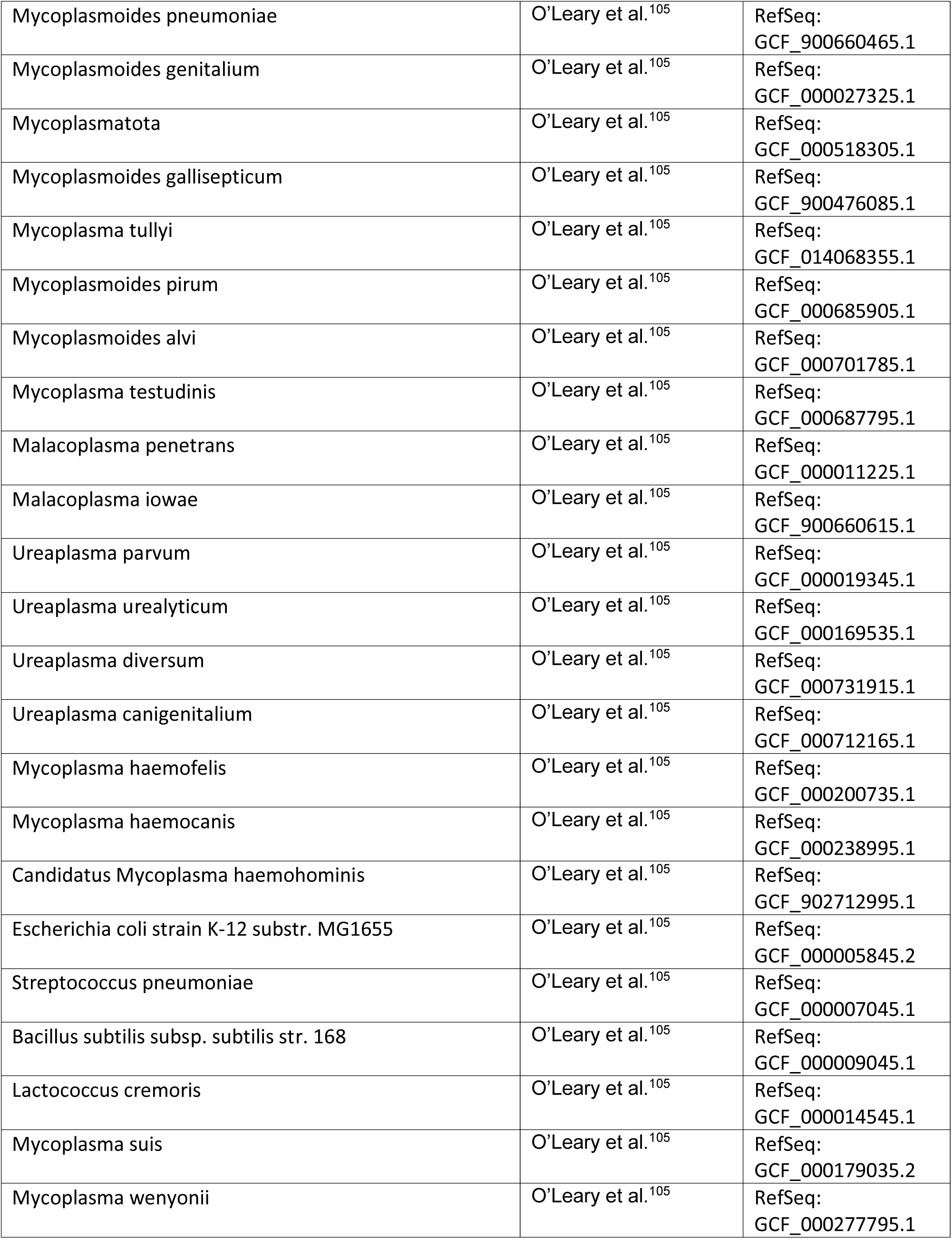
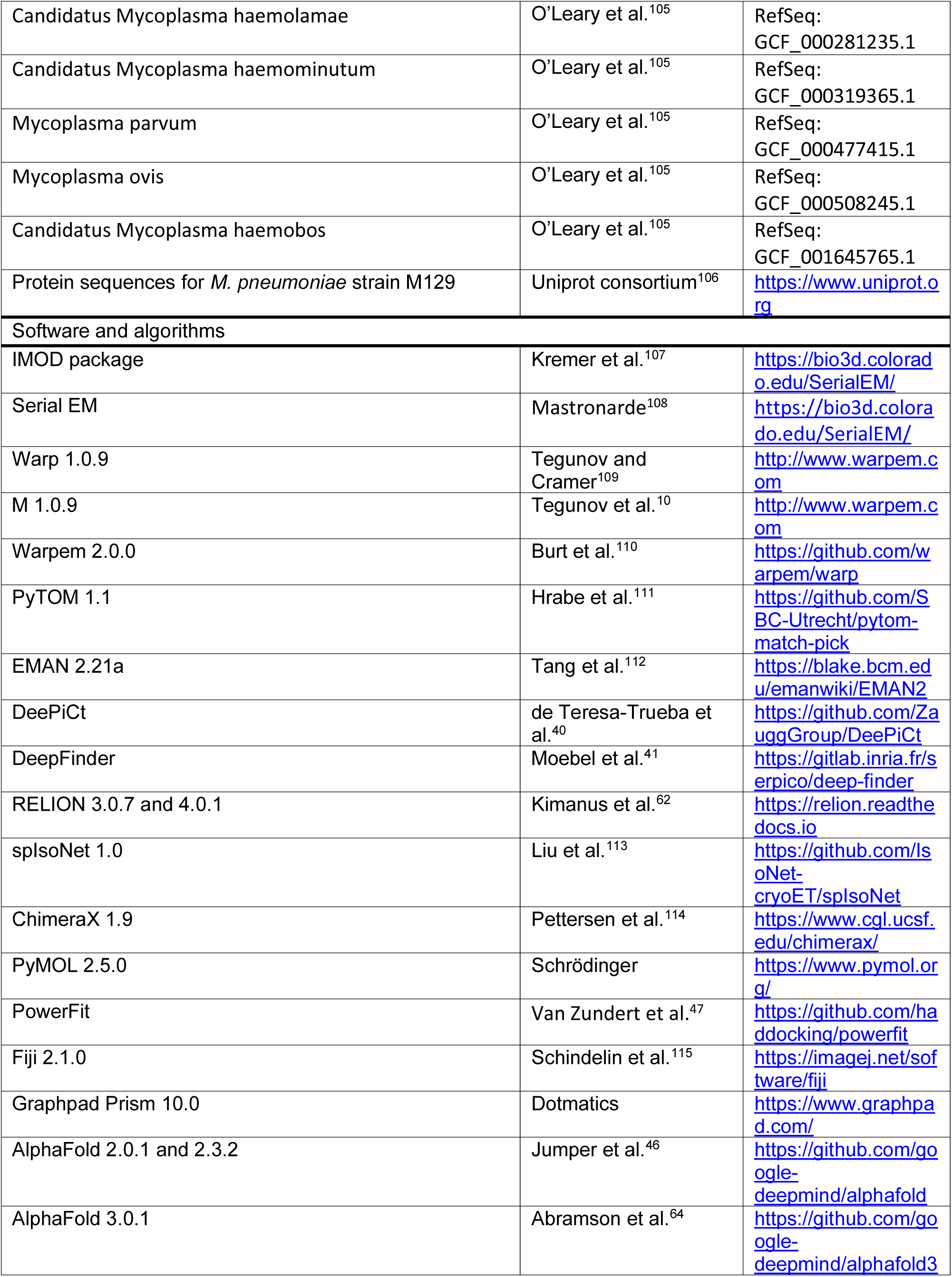
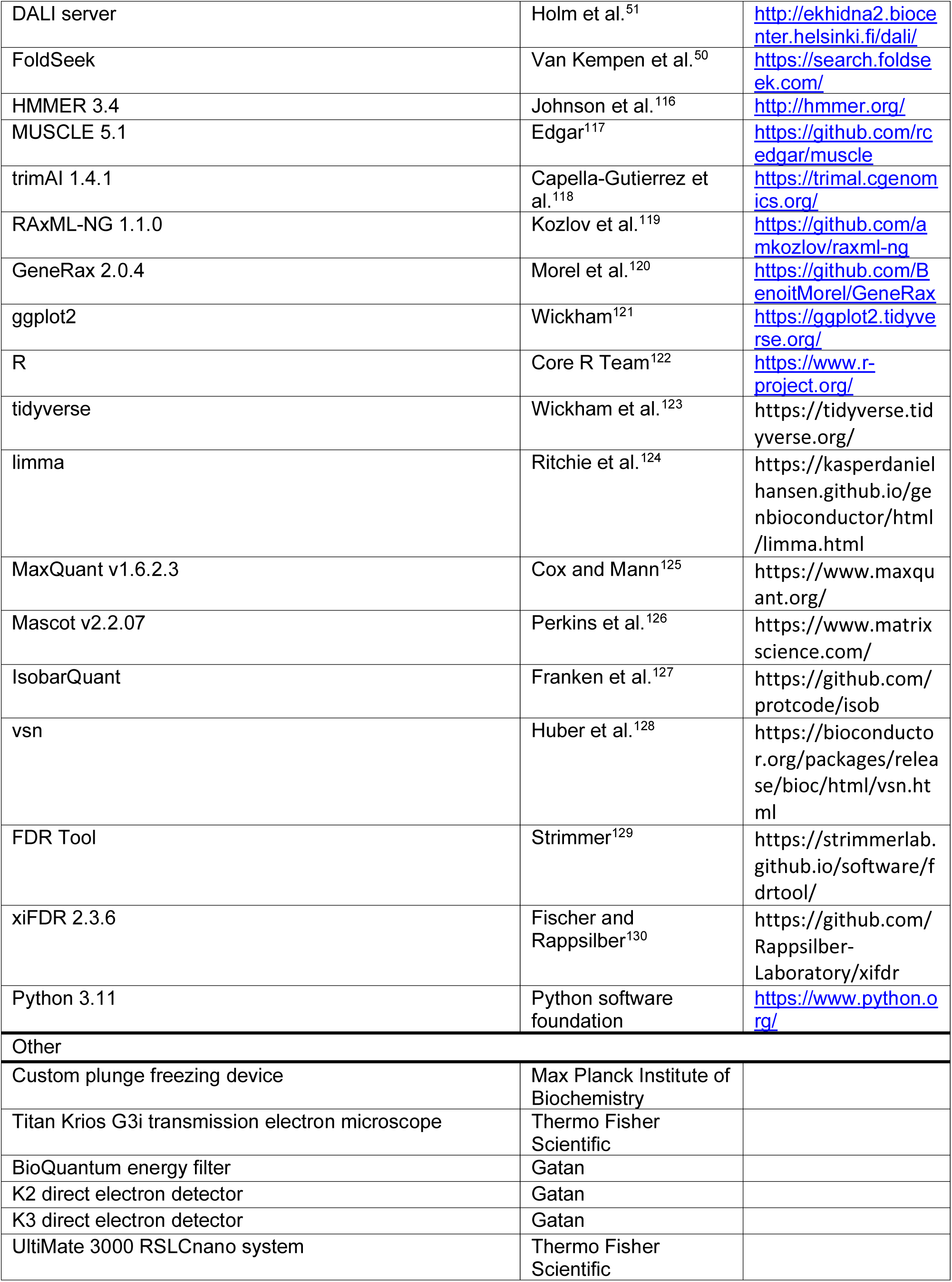
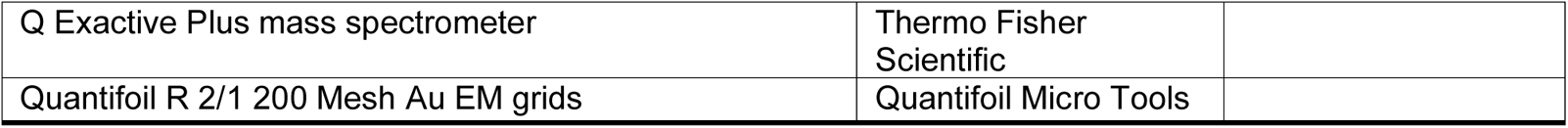

### Resource availability

#### Lead contact

Further information and requests for resources and reagents should be directed to and will be fulfilled by the lead contact, Julia Mahamid.

## Materials availability

This study did not generate new reagents

## Data and code availability

- Cryo-ET maps will be deposited at the Electron Microscopy Databank (EMDB). The models will be deposited in Model Archive. The maps and models will be made public upon publication, with their accession codes listed in the key resources table.
- Structures for initial model building were obtained from the Protein Data Bank (PDB) with accession codes 8P8B and 8P6P or generated with AlphaFold2. Structures used for comparison with the models generated in this work are listed with accession codes in the key resources table.
- This study does not report any original software packages or code.
- Any additional information required to reanalyze the data reported in this work is available from the lead contact upon request.

## Experimental model

*Mycoplasma pneumoniae* M129 (ATCC 29342) cells were kindly provided by Professor Jörg Stülke. Cell cultivation was done as previously described^17,20^ in modified Hayflick medium at 37°C.

## METHOD DETAILS

### Cryo-ET data collection and initial processing

*Mycoplasma pneumoniae* sample preparation and cryo-ET data collection were performed as previously described^17,20^. In brief, *M. pneumoniae* cells were grown on Quantifoild gold grids coated with R2/1 holey carbon film and vitrified using a manual plunger. Tilt-series data were collected on a Titan Krios G3 microscope equipped with a Gatan direct electron detector following the dose-symmetric scheme^131^ (tilt angles -60°:60° with 3° step and total dose 120-150 e^−^/Å^2^). Five datasets were analysed in this study, which differ in detector type (K2 or K3), data pixel size, and drug treatment (Table S1). For the untreated cells dataset (here referred as dataset I), 355 tilt-series were collected (K2 camera, magnification 64k, pixel size 1.7 Å). For the four datasets of antibiotic-treated cells (here referred to as datasets II, III, IV and V), saturating concentrations of the different drugs were added to the culture dishes and the treatment time was controlled to be 15 to 20 minutes. Details for dataset II (65 tilt-series of chloramphenicol treated cells, K2), dataset III (86 pseudouridimycin treated cells, K2), dataset IV (70 spectinomycin treated cells, K3), dataset V (139 chloramphenicol treated cells, K3) can be found in previous publications^17,20,61^.

Tilt-series processing was performed in either Warp versions 1.0.7 or 1.0.9^10,109^. Ribosome-based multi-particle refinement of the tilt-series, including both geometric and CTF parameters, was done in the software M^10^ associated with Warp. All subtomograms used for structure determination were reconstructed from tilt-series after M refinement unless otherwise indicated. Dataset I was additionally processed in Warp-2 (warp-2.0.0dev31)^110^, using default parameters unless otherwise indicated. Initially, the CTF was re-estimated using WarpTools fs_ctf on a 2x2 grid to a maximum resolution of 10 Å, with a max defocus value of 6 µm. The previous tilt-series alignments were read into Warp using WarpTools ts_import and WarpTools ts_import_alignments, where 3D CTF estimation was performed with WarpTools ts_ctf using the same parameters as for 2D CTF estimation. The tilt series were subsequently refined using MCore based on the ribosome poses in an iterative manner. In all rounds, default parameters were used and particle orientations were refined unless otherwise indicated. First, the tilt alignments were corrected for gross errors using a 1x1 image warp grid. Subsequently, a round of refinement was performed using a 2x2 image grid while refining stage angles, resulting in a ribosome map with resolution of 5.2 Å. Next, local defocus was refined using an exhaustive search and a 2x2 image warping grid without refinement of particle orientations. This was followed by a round of volume warping on a 2x2x2x10 grid and a round of image warping on a 2x2 grid and local defocus refinement, resulting in a resolution of 3.7 Å. This was used as input for a round of global tilt weighting followed by tilt-series weighting using EstimateWeights. The output was then used for a final round of refinement in MCore using a 2x2 image warping grid and another round of local defocus refinement, resulting in a final ribosome map with a resolution of 3.5 Å.

### Particle picking of the ribosome

Ribosome positions in all datasets, localized based on template matching with PyTom version 1.1^111^ and RELION classification as the final cleaning procedure, were reused from previous work^10,17,20^. In total, 77,539 untreated (dataset I), 18.987 Cm-treated (dataset II), 15,314 PUM-treated (dataset III), 12,935 spectinomycin-treated (dataset IV), and 30,774 Cm-treated (dataset V) ribosome subtomograms were reconstructed using Warp for further analysis.

### Particle picking and initial subtomogram averaging of the dome complex

Initially, 1000 dome complexes from 50 tomograms were manually picked in the untreated data (dataset I) using EMAN2.21a e2spt_boxer.py^112^. Subtomograms were reconstructed in Warp, and initial subtomogram averaging was performed in RELION-4.0.1^62^. The refined average, coordinates, and particle orientations were used to create a binary mask for each tomogram using a python script. The binary segmentations were used as training data for DeePiCt^40^ to pick additional particles in the tomograms. The DeePiCt model was trained on 6.8 Å/pixel tomograms over 200 epochs using default parameters, except for depth of 3, initial features of 16, box size of 64^3^ voxels and overlap of 12 voxels. After training and prediction, post processing was performed with a threshold of 0.985 and minimum cluster size of 3500 voxels. The resulting coordinates were used to reconstruct subtomograms in Warp and for subtomogram averaging and 3D classification in RELION to sort out true/false positives. The training, prediction and curation were done iteratively for four rounds to obtain the most complete and accurate particle positions. Subsequently, this list of localized dome complexes was used in concert with the ribosome positions to train a multiclass DeepFinder network^41^. To avoid particle picks above and below the cells, 200 spheres with a diameter of 150 Å were randomly inserted in the upper and lower 50 slices of the tomogram, and added as an additional background class for training. This significantly improved localization accuracy. DeepFinder was trained on 6.8 Å/pixel tomograms with default parameters, except for a patch size of 60^3^ voxels for 200 epochs. This was then used to localize particles in all tomograms. The outputted labels were binned to 13.6 Å/pixel, and post processed with a threshold size of 300 pixels and cluster radius of 6 pixels. Finally, the positions were manually curated using EMAN2.21a e2spt_boxer.py resulting in 13,770 particles in the untreated data over 355 tomograms, and 6,103 particles in the antibiotics-treated dataset over 183 tomograms (Table S1).

### Subtomogram analysis of the major closed and SecA minor conformations of the dome

Subtomograms were reconstructed at 6.8 Å/pixel at a box size of 64^3^ voxels in Warp 1.0.9 and processed as described in Figure S1B. First, an initial model was generated in RELION version 4.0.1 using the 3D initial model algorithm, followed by a 3D refinement with global searches using a diameter of 220 Å for both normalization and a spherical mask. This was used for the first 3D classification (I), where a 50 Å spherical mask focused on SecA was used without alignment. The classification was performed four times using T values of 0.1, 1, 5, and 20, which all yielded similar results. The classifications separated particles to the major (closed, 11,557 particles) and the minor SecA (2,213 particles) conformations. Particles were then exported with WarpTools ts_export_particles in 3D mode with a pixel size of 3.4 Å/pixel, box size of 112^3^ voxels, and a normalization diameter of 220 Å. A 3D refinement was performed for both classes and the resulting maps were used to generate loose particle-shaped masks. The masks were used in a final 3D refinement with local angular searches, resulting in 13 Å and 9.5 Å global resolution maps for the SecA minor conformation and the major closed conformation, respectively. The particles corresponding to the closed conformation were imported into M in Warp-2.0, and used for a round of refinement of particle poses, followed by two rounds of both particle pose and image warping (2x2 grid) refinement. This resulted in a map with a global resolution of 9.12 Å, which was used for one round of defocus refinement, and volume warping (2x2x2x10 grid), resulting in a 9.1 Å resolution. The data were then resampled to 1.7 Å/pixel, followed by two M refinements of image warping (2x2 grid) and defocus, resulting in the final map with a global resolution of 9.07 Å.

### Subtomogram averaging of additional states of the dome complex

The particles from the closed conformation were further classified in RELION as shown in Figure S1B. First, the particles were refined once using spIsoNet^113^, and used for a second 3D classification (II) without alignment focused on SecDF using a 60 Å spherical mask. For all classification, unless otherwise stated, T values of 0.1, 1, 5, and 20, and a class number of 5 were used. All classifications, except for T=20, resulted in clear separation of 3 distinct populations, corresponding to the original closed conformation (major class), an Mdp444 minor conformation (2,010 particles), and a conformation where the SecDF was rotated inward (1,459 particles). The Mdp444 minor conformation was subjected to another round of 3D classification focused on SecDF, which revealed a high level of flexibility in the interaction between SecDF and Mdp444. The major class was used as a reference for a final round of local 3D refinement, resulting in the final map at a global resolution of 14 Å. The alternative SecDF conformation (1,459 particles) was subjected to another round of SecDF focused classification (III) without alignment, revealing two distinct subpopulations of particles. One corresponded to an inward-facing SecDF (1,047 particles), which still associated with Mdp444, and another corresponded to a more open conformation (412 particles) that no longer interacted with Mdp444. Classification of the inward-facing SecDF (1,047 particles) with a 50 Å spherical mask focused on the dome-interior (IV; substrate mask), further separated the inward-facing SecDF conformation into substrate-bound (588 particles) and substrate-less (459 particles) classes. Global refinement of both the inward-facing SecDF conformation and the inward-facing SecDF with substrate conformation resolved the maps to a resolution of 25 Å. The more open class (412 particles) from classification III, was subjected to a final focused classification on SecA (V), which revealed both a SecA-bound (235 particles) and a SecA-less (177 particles) open conformations. The two classes were subjected to a final 3D refinement with global searches, resulting in global resolutions of 27 Å and 29 Å for the open and open with SecA conformations, respectively.

### Spatial analysis of the dome complex

Number of particles per tomogram, the mean and SD (39±17), were calculated and reported in Figure 1B. For investigating dome clustering, center coordinates with respect to the tomogram frame of reference for each particle were calculated by applying the translational shifts determined during 3D refinement to the original particle coordinates. Subsequently, the pairwise distances between all particles within each tomogram were calculated, and particles closer than approximately 1.3 times the particle diameter (270 Å) were designated as being in a cluster. Finally, any clusters with overlapping particles were merged. The number of particles per cluster was calculated and plotted (Figure 1C) in GraphPad Prism Version 10.0 (Dotmatics).

For investigating the spatial association between the dome complex and the ribosome, we first defined the positions corresponding to the SecA PPXD domain in the dome complex and the ribosome nascent peptide exit site with respect to their centers using ChimeraX version 1.9 (USCF), and validated them in IMOD^107^. The coordinates corresponding to these positions in the tomogram frame of reference were updated for each particle, using the original coordinates, euler angles, and translational shifts determined during 3D refinements. Finally, the nearest ribosome peptide exit site was determined for each dome by calculating the distance to all peptide exit sites and taking the smallest value. The histogram of distances was then calculated and plotted in GraphPad Prism version 10.0 (Dotmatics) as shown in Figure 5B.

### Subtomogram analysis of the ribosome-dome supercomplex

Ribosome subtomograms were reconstructed with a large box size (voxel size 2.9 Å, 256^3^ voxels) to accommodate potential neighbouring dome complexes, and subjected to classification with a mask focused on the peptide exit site region in RELION 3.0.7 as shown in Figure S5. For the native untreated dataset (I), this classification of the total 77,539 ribosome subtomograms did not result in maps representing ribosome-dome complexes, likely due to the low frequency of bound dome complexes. For this dataset, we therefore instead first filtered-out ribosomes for which the peptide exit tunnel was positioned at distance of more than 20 nm from their closest dome neighbour, resulting in only 1,791 ribosomes. Focused classification for this subset of ribosomes resulted in 151 closely assembled ribosome-dome complexes, which were subsequently subjected to refinement. For the four datasets of antibiotics treatments (II, III, IV, V), exhaustive focused classification^20^, detailed in Figure S5G, was performed with at least three RELION jobs. In total, 2,304 supercomplexes were pooled from the four datasets and subjected to refinement. Finally, focused refinement, multi-body refinement, local resolution estimation and post-processing were done in RELION.

### Surface shaving sample preparation and measurement

*M. pneumoniae* cells were seeded into a T75 EasYFlask (Nunc) and grown for four days at 37°C as described above. The medium was removed and the cells were washed twice in PBS. The cells were then detached using a cell scraper, washed twice with PBS by pelleting the cells at 1,000 g for 10 minutes in a centrifuge cooled to 4°C, and resuspended in PBS. The cell density was measured at 600 nm using a MicroSpek DSM Dual Cell density photometer (Laxco Inc) and the cells were diluted to OD600=1.11. Subsequently, either trypsin (Promega; 2 µg/mL), proteinase K (Promega; 500 µg/mL) or PBS (control) were added to the cell suspension to a final OD600=1. The cells were incubated for 15 minutes at 37°C shaking, and the reaction was stopped by addition of 2.5 mM phenylmethylsulfonyl fluoride. The cells were then recovered by centrifugation for 10 minutes at 1,000 g in a centrifuge cooled to 4°C. Cell pellets were resuspended in 250 µL of lysis buffer (2% SDS in LC-MS/MS H_2_O), incubated on ice for 5 minutes, then centrifuged at 20,000 g to remove insoluble debris. Supernatants containing cellular protein extracts were transferred to a fresh microfuge tube. Proteins were digested according to a modified SP3 protocol ^132,133^. Briefly, approximately 5 μg of protein was diluted in buffer containing a final concentration of 1.65% SDS and 16.6 mM Tris(2-carboxyethyl)phosphine hydrochloride (TCEP) in a 20 µL volume. Samples were added to a bead suspension (10 μg of beads (Thermo Fischer Scientific—Sera-Mag Speed Beads, (4515-2105-050250 and 6515-2105-050250) in 10 μl 15% formic acid plus 30 μl ethanol). After a 15-min incubation at room temperature with shaking at 500 rpm, beads were washed four times with 70% ethanol. Next, proteins were digested overnight by adding 40 μl of digest solution (5 mM chloroacetamide, 1.25 mM TCEP, 200 ng trypsin and 200 ng LysC in 100 mM HEPES pH 8). Peptides were then eluted from the beads, dried under vacuum, reconstituted in 10 μl of water and labelled with shaking at 500 rpm for 30 min at room temperature with 45 μg of TMT10plex^TM^ (Thermo Fisher Scientific) dissolved in 4 μl of acetonitrile. The reaction was quenched with 4 μl of 5% hydroxylamine for 15 min at room temperature with shaking at 500 rpm, and experiments belonging to the same mass spectrometry run were combined into one sample. Samples were desalted with solid-phase extraction by loading the samples onto a Waters OASIS HLB μElution Plate (30 μm), washing them twice with 100 μl of 0.05% formic acid, eluting them with 100 μl of 80% acetonitrile and 0.05% formic acid, and drying them under vacuum. Finally, samples were fractionated into 6 fractions on a reversed-phase C18 system running under high pH conditions. This consisted of an 85 min gradient (mobile phase A: 20 mM ammonium formate (pH 10) and mobile phase B: acetonitrile) at a 0.1 ml/min starting at 0% B, followed by a linear increase to 35% B from 2 to 60 min, with a subsequent increase to 85% B from up to 62 min and holding this up to 68 min, which was followed by a linear decrease to 0% B up to 70 min, finishing with a hold at this level until the end of the run. Fractions were collected every 2 min from 12 to 70 min and every 6^th^ fraction was pooled together.

Samples were analyzed by liquid chromatography coupled to tandem mass spectrometry, as previously described^134^. Briefly, peptides were separated using an UltiMate 3000 RSLCnano system (Thermo Fisher Scientific) equipped with a trapping cartridge (Precolumn; C18 PepMap 100, 5 μm, 300 μm i.d. × 5 mm, 100 Å) and an analytical column (Waters nanoEase HSS C18 T3, 75 μm × 25 cm, 1.8 μm, 100 Å). Solvent A was 0.1% formic acid in LC-MS grade water, and solvent B was 0.1% formic acid in LC-MS grade acetonitrile. Peptides were loaded onto the trapping cartridge (30 μl/min of 0.05% trifluoroacetic acid in LC-MS grade water for 3 min) and eluted with a constant flow of 0.3 μl/min using a 120 min analysis time (with a 2–30% B elution, followed by an increase to 40% B, and a final wash to 80% B for 2 min before re-equilibration to initial conditions). The LC system was directly coupled to a Q Exactive Plus mass spectrometer (Thermo Fisher Scientific) using a Nanospray-Flex ion source and a Pico-Tip Emitter 360 μm OD × 20 μm ID; 10 μm tip (New Objective). The mass spectrometer was operated in positive ion mode with a spray voltage of 2.3 kV and capillary temperature of 275°C. Full-scan MS spectra with a mass range of 375–1,200 *m*/*z* were acquired in profile mode using a resolution of 70,000 (maximum fill time of 250 ms or a maximum of 3e6 ions (automatic gain control, AGC)). Fragmentation was triggered for the top 10 peaks with charge 2–4 on the MS scan (data-dependent acquisition) with a 30-s dynamic exclusion window (normalized collision energy was 30), and MS/MS spectra were acquired in profile mode with a resolution of 35,000 (maximum fill time of 120 ms or an AGC target of 2e5 ions).

### Surface shaving mass spectrometry data processing

MS data were processed as previously described using Maxquant version 1.6.14.0^127^. MS/MS spectra were searched against the *Mycoplasma pneumoniae* proteome (UniProt Proteome ID: UP000000808), supplemented with common contaminants and reversed sequences for False Discovery Rate (FDR) estimation. The search was conducted using Trypsin/P and LysC specificity with up to two missed cleavages allowed. Carbamidomethylation of cysteine was set as a fixed modification; oxidation of methionine and N-terminal acetylation were used as variable modifications. The peptide and protein FDRs were controlled at 1%. Reporter ion MS2-based TMT quantification was used with a minimum reporter ion count of 2. A minimum peptide length of 7 amino acids was required. Modified peptides required a minimum score of 40 and delta score of 6 for identification. Only unique peptides were used for quantification, and the “match between runs” feature was disabled. For peptide-level abundance score calculations, peptide and protein intensities were obtained from MaxQuant output files (peptides.txt and proteinGroups.txt). Peptides annotated as reverse hits or potential contaminants were excluded. The reporter ion intensities (TMT10plex) were log2-transformed and normalized using the vsn package in R to account for variation across TMT channels. To correct for inter-channel intensity differences, log2 intensities were median-centered per channel using the overall distribution across all peptides. Abundance scores for each peptide were computed by creating a unique identifier by combining the protein accession with its start and end positions. The median normalized intensity across replicates was computed for each experimental condition. Abundance scores were defined as the log2 fold change in peptide intensity between protease-treated samples (either trypsin or proteinase K) and corresponding untreated controls, calculated across three biological replicates. Statistical testing and multiple testing correction was performed as follows. Linear modeling was performed separately for each protease comparison using the limma package. The model included treatment (protease vs. control) and replicate effects. Moderated t-statistics and p-values were extracted, and FDRs were computed using the fdrtool package. Peptides with FDR-adjusted p-values below 0.05 were considered significantly different in abundance. Peptides were annotated using sequence headers from the *M. pneumoniae* UniProt reference proteome. Gene names, protein descriptions, and amino acid positions were extracted and merged with statistical results. Volcano plots were generated using ggplot2 and ggrepel to visualize significant abundance changes.

### Re-analysis of crosslinking MS data

DSSO search results of a previously published *M. pneumoniae* crosslinking MS dataset^17^ (Proteome exchange database ID: PXD017711) were re-analyzed. Prior to FDR calculation, matches were filtered for having at least one peptide with a DSSO fragment, 4 matched fragments per peptide and a delta score of 33% of the match score. Results were filtered for an FDR of 5% on residue pair level and 10% on protein-protein interaction level using xiFDR version 2.3.6.

### AlphaFold predictions

AlphaFold v2.0.1^46^ was used to predict the structure of the 113 identified *M. pneumoniae* surface proteins. Default parameters were used, except for number of recycles set to 5, and only one model being generated for each of the five weight sets (num_prediction_per_model=1). The top ranked model was selected for each protein. Subsequently, the structure of the Mdps, SecDF, SecA, MPN523, and the SecYEG complex were predicted using AlphaFold v2.3.2 multimer^46,135^ with 30 recycles and five models generated for each weight set. The AlphaFold v3.0.1^64^ of SecYEG in complex with SecDF was predicted using three diffusion samples and otherwise default parameters.

### Rigid-body fitting of AlphaFold predictions of *M. pneumoniae* surface proteins

The cryo-ET map of the closed conformation was masked to only include the extracellular region of the complex using a cylindrical mask in ChimeraX 1.9 (UCSF). Subsequently, the predicted structural models were individually rigid-body fitted against the map using PowerFit^47^, at a resolution of 15 Å and a rotational sampling of 7°. The resulting cross-correlation values were plotted as a function of the respective proteins molecular weight to identify Mdp436, Mdp444, and Mdp489 as candidates for the dome complex (Figure 2B and Table S3).

### Structural modeling of the dome complexes

The cryo-ET map of the closed conformation was used for rigid-body fitting of the AlphaFold2 monomer models of the *M. pneumoniae* MPN523, the Mdps, and SecA in PowerFit^47^ at a resolution of 10 Å. The same procedure was used for the AlphaFold2 multimer model of the SecYEG complex. By visually inspecting the fits, regions that were deemed to not fit well, were refined using ChimeraX version 1.9. The AlphaFold2 monomer model of SecDF was separated into the extracellular (residues 45-501) and transmembrane (residues 1-38 and 514-927) regions, which were separately fitted to the map using fit-in-map in ChimeraX.

For the open conformation, this model of MPN523, the Mdps and SecA was initially fitted into the open conformation cryo-ET map derived from antibiotics-treated cells. The transmembrane regions of SecDF and SecYEG were reorganized compared to the closed conformation (Figures 6E-F and S7B-D), and we therefore used AlphaFold 3^64^ to predict the SecDF-SecYEG subcomplex. This model was fitted as a single rigid body using ChimeraX. Mdp444 was substantially tilted, compared to the closed conformation, and was therefore fitted separately to the map using ChimeraX. As density for SecA was not present in the map, it was removed from the model.

For the ribosome-dome supercomplex, the atomic models of the 50S and 30S ribosomal subunits from *M. pneumoniae* (pdb: 8P8B and 8P6P) were first rigid-body fitted into the consensus refinement using Chimera and the remaining densities were segmented and modelled in a sequential manner separately from the procedure described above. The AlphaFold2 predicated models for the whole proteome of *M. pneumoniae* strain M129 (ATCC 29342) were downloaded from the AlphaFold Database (https://alphafold.ebi.ac.uk, accessed during July 2022). Systematic fitting of models into the remaining densities was done using Chimera and Assembline scripts^136^. For the extracellular dome density, MPN444, MPN489 and MPN436 provided high fitting scores and good fitting quality under visual inspection, which aligned with the results of the systematic fitting of the standalone dome complex. Subsequently, guided by the crosslinking MS data, MPN523 and SecDF were placed into the map by rigid-body fitting, and explained the rest of the density. For the intracellular density, the region between the ribosome and the plasma membrane was unambiguously fitted with SecA due to the relatively high local resolution of about 8 Å, enabling confident modeling at the secondary structure level. After fitting all the above proteins, the remaining channel-like density across the membrane beneath SecA resembled the previously reported SecA-SecE structures^36,37,59,60^. Fitting of the model of SecA and SecYEG (MPN184, MPN068, MPN242) predicated by AlphaFold2 multimer^135^ explained the transmembrane density. In the final model, no major density remained unassigned, except for the nascent peptide density in the ribosome exit tunnel and inside the cavity of the dome complex.

### Domain rotation and distance measurements in PyMOL

All SecY-SecA structures (pdb: 3DIN, 6ITC, 5EUL, 7XHA, and 7XHB) were aligned on the Cα atoms of SecY in the closed dome complex model, using the ’super’ command in PyMOL version 2.5.0 (Schrödinger). The angle and translational parameters of the position of SecA, as well as the RMSD values, were then calculated using the script https://pymolwiki.org/index.php/Angle_between_domains as part of the psico script package (https://github.com/speleo3/pymol-psico). Similarly, for SecDF, the open conformation model was aligned to the closed conformation model of the dome complex, based on the Mdp489 and Mdp436 coordinates. Subsequently, the rotation angle was determined between the two extracellular domains of SecDF (head and base, residues 45-501) in the open and closed conformation, respectively, using the same script. The distance measurements for the opening of the dome in the closed, open and wide-open conformation were performed by rigid-body fitting the Mdp444 and Mdp489 AlphaFold2 derived models into each map, and measuring the distance from the Cα atoms of Mdp444 residue 1124 to Mdp489 residue 178 using the distance command in PyMOL version 2.5.0.

### Model preparation, DALI and FoldSeek structural homology searches

The predicted AlphaFold structures of the Mdps were aligned in PyMOL version 2.5.0 (Schrödinger), and aligned on their Cα carbons using the ’super’ command. Subsequently, the domain boundaries were determined by manual inspection, resulting in 6 domains for each Mdp. The domains were defined as follows: For Mdp436, domain 1: residues 54-159, and 777-811; domain 2: residues 159-222, and 574-627; domain 3: residues 224-233, 273-285,306-360, 471-478, and 515-536; domain 4: residues 818-950, and 1179-1244; domain 5: residues 955-962, 1037-1063, and 1162-1179; domain 6: residues 1065-1143. For Mdp444, domain 1: residues 53-160, and 719-759; domain 2: residues 161-228, and 511-570; domain 3: residues 229-239, 277-369, and 466-510; domain 4: residues 759-844, and 1273-1325; domain 5: residues 903-948, 1035-1059, and 1211-1245; domain 6: residues 1105-1167. For Mdp489, domain 1: residues 56-166, and 817-855; domain 2: residues 156-222, 543-605, and 641-650; domain 3: residues 222-233, 281-293, 328-384, 455-462, and 515-531; domain 4: residues 856-915, 957-1007, and 1286-1300; domain 5: residues 1012-1020, 1119-1149, 1235-1245, and 1272-1278; domain 6: residues 1163-1232. For MPN523, we removed the signal peptide, the long disordered loop, and the loop region extending toward Mdp436, leaving residues 30-147, 202-236, and 292-304. To search for homologs of each domain or protein, we used the FoldSeek server^50^, searching against the AlphaFold/Uniprot50 database. Two separate runs were performed for each domain using either mode 3Di/AA or TM-align. All results are reported in Table S4. For the DALI server^51^, we ran each domain or protein against the full PDB. All results are reported in Table S5.

### Phylogenetic analysis of Mdp436, Mdp444, and Mdp489

The phylogenetic analysis involved two steps: reconstruction of the species tree, and reconciliation of the gene tree with the species tree using the duplication/loss model. First, we selected all the proteins homologous to the Mdps by performing a hhblits (version 3.3.0) search^137^ against the RefSeq database^105^ with default parameters. We noted that all of the significant hits belonged to species in the *Mycoplasmoidaceae* family. When two or more strains from the same species were reported, only the type strain was retained, except for *M. pneumoniae*, where the strain M129 used for our experiments was kept instead of the type strain FH. The representative genomes for all the species in the *Mycoplasmoidaceae* family were downloaded from RefSeq using the "dataset" command-line utility [https://www.ncbi.nlm.nih.gov/datasets/docs/v2/reference-docs/command-line/datasets/]. To the proteins that were reported by hhblits, we manually added PrsA which we identified from the structural homology searches. PrsA was added from the model organisms, *E. coli*, *B. subtilis*, *L. cremoris*, and *S. pneumoniae*, in addition to their genomes downloaded from RefSeq. We used GTDB-tk v2^138^ to extract the default 120 marker genes from these species (using the "identify" command) and perform a multiple sequence alignment (using the "align" command). We then reconstructed the species tree using FastTree 2^139^, and rooted it using *E. coli* as the outgroup.

For the gene tree reconciliation, the protein sequences of the hits from hhblits and the PrsA sequences from the four selected species were concatenated in a FASTA file and aligned with MAFFT v7.520^140^. Since there were proteins of diverse lengths, ranging from 150 to 1365 amino acids, the alignment was performed in two steps: first, only the proteins longer than 600 amino acids were aligned using the "auto" strategy; then, the remaining, short sequences were added to the alignment using the "--addfragments" and "--keeplength" options of MAFFT. The alignment was trimmed with trimAL v1.4.1^118^ using the "automated1" option, and the trimmed alignment was finally processed with GeneRax^120^ together with the species tree built in the previous step. GeneRax performed the reconciliation using the "UndatedDL" model and produced an output file in RecPhyloXML format^141^, which was visualized using the ggplot2 package^121^ in the R programming language^122^.

### Phylogenetic analysis of MPN523

We used the DALI server^51^ to perform a pairwise structure alignment between the AlphaFold-predicted structures of MPN263 (thioredoxin A) and MPN523. As mentioned above, before executing the alignment, we excised signal peptide and disordered loops from the AlphaFold2 prediction of MPN523 (Residues retained: 30-147, 202-236, 292-305). The aligned residues had 14% sequence identity, and the pairwise structural alignment had a Z-score of 5.0 and a root mean square deviation of 3.1. We extracted the sequence alignment from the DALI results and built a corresponding hidden Markov model (HMM) profile using HMMER v3.4^116^. Such HMM captures sequence peculiarities of both MPN523 and of the more classic thioredoxin MPN263. Relying on the sequence database of 28 species that we previously built, including the whole *Mycoplasma* family and its closest well-studied relatives (*E. coli*, *B. subtilis*, *S. pneumoniae*, and *L. cremoris*), we performed four iterations of ’hmmsearch’, using the output of the previous iteration to build the HMM for the next round. This yielded a set of 78 protein sequences from the 28 species that are homologous to both MPN523 and MPN263. We aligned the sequences using MUSCLE v5.1^117^ with the ’super5’ algorithm and trimmed the alignment using trimAl v1.4.1^118^ with the ’automated1’ option. We built a tree out of the trimmed alignment with RAxML-NG v1.1.0^119^, using the ’LG+G8+F’ model and starting from 10 parsimony trees. Finally, to disentangle the orthologs of MPN523 from the paralogs, we also built a reconciliated gene/species tree starting from the same trimmed alignment. To do this, we used GeneRax v2.0.4^120^ with the ’LG+G’ substitution model and ’UndatedDL’ reconciliation model. The phylogenetic tree of the lineage that is orthologous to MPN523, was visualized using the ggplot2 package^121^ in the R programming language^122^.

## QUANTIFICATION AND STATISTICAL ANALYSIS

The details of the quantification and all statistical analyses are included in figure captions or the relevant sections of method details.

