## Supplemental information for "In-cell discovery and characterization of a non-canonical bacterial protein translocation-folding complex"

### Supplementary information

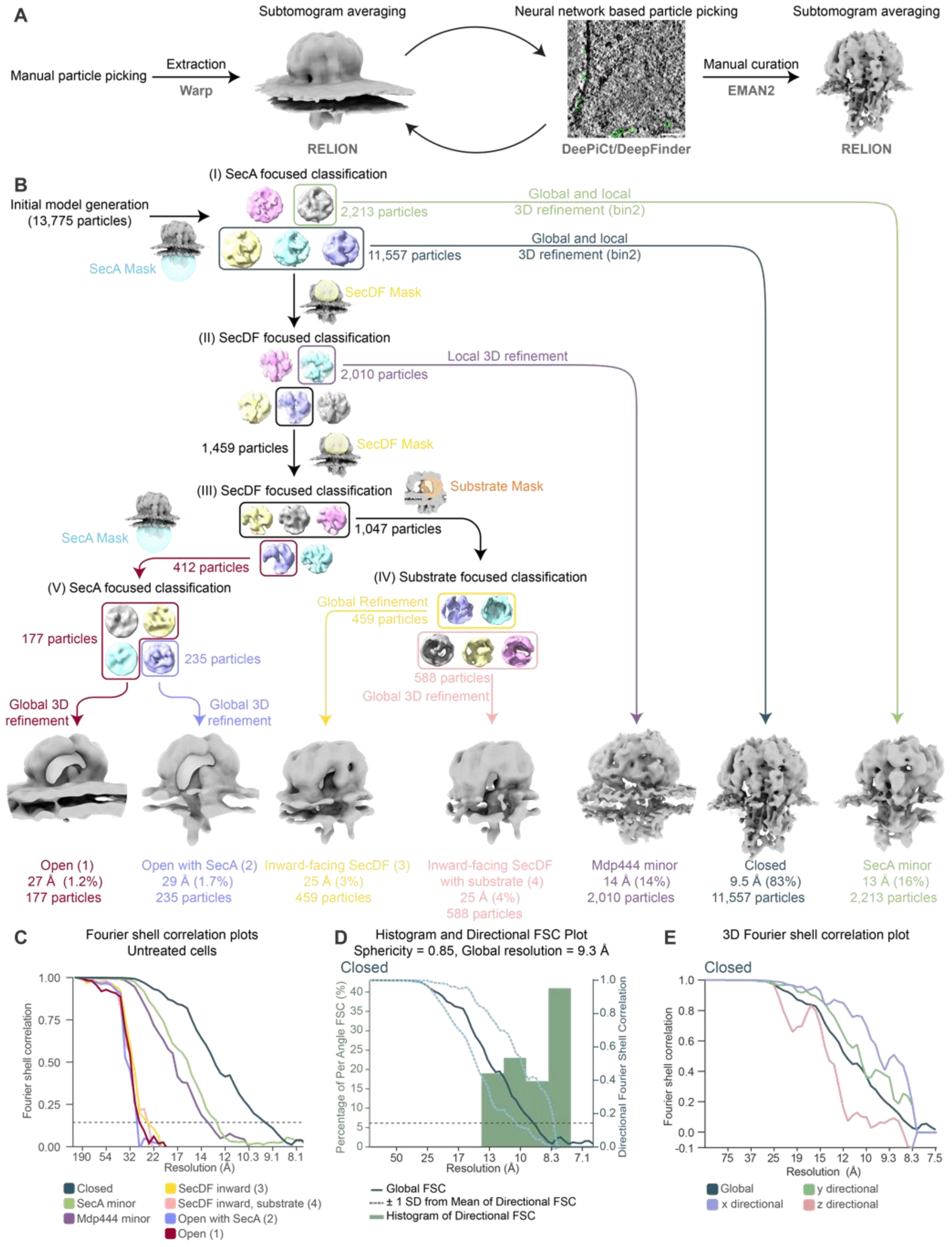


**Supplementary Figure 1. Workflow for particle picking and subtomogram analysis of the dome complex in untreated *Mycoplasma pneumoniae* cells.** Related to Figure 1.

(A) Particle picking pipeline. Particles were initially manually picked, and subtomograms reconstructed with Warp. Subtomogram averaging and classification were performed in RELION, and the outputted map, particle positions and Euler angles were used to create a 3D segmentation for each tomogram. The segmentations were subsequently used to train a DeePiCt or DeepFinder model for picking of additional particles. The RELION-DeePiCt/DeepFinder pipeline was performed iteratively to obtain the most complete and accurate particle picks. Each particle position was manually validated, and missed particles were added, leading to the final particle list used for downstream structural analyses. (B) Pipeline for subtomogram averaging and hierarchical classification in RELION. The different classification and refinement strategies are indicated for each arrow. Masks used for classification are indicated at each level. Particle counts and global resolution are shown for each of the final maps. (C) Fourier shell correlation (FSC) plots for the maps shown in B. FSC threshold 0.143 used for estimating global resolution is shown as dashed line. (D-E) 3D FSC plots for the major closed conformation shown in B, indicate that the map is slightly anisotropic (sphericity of 0.85).

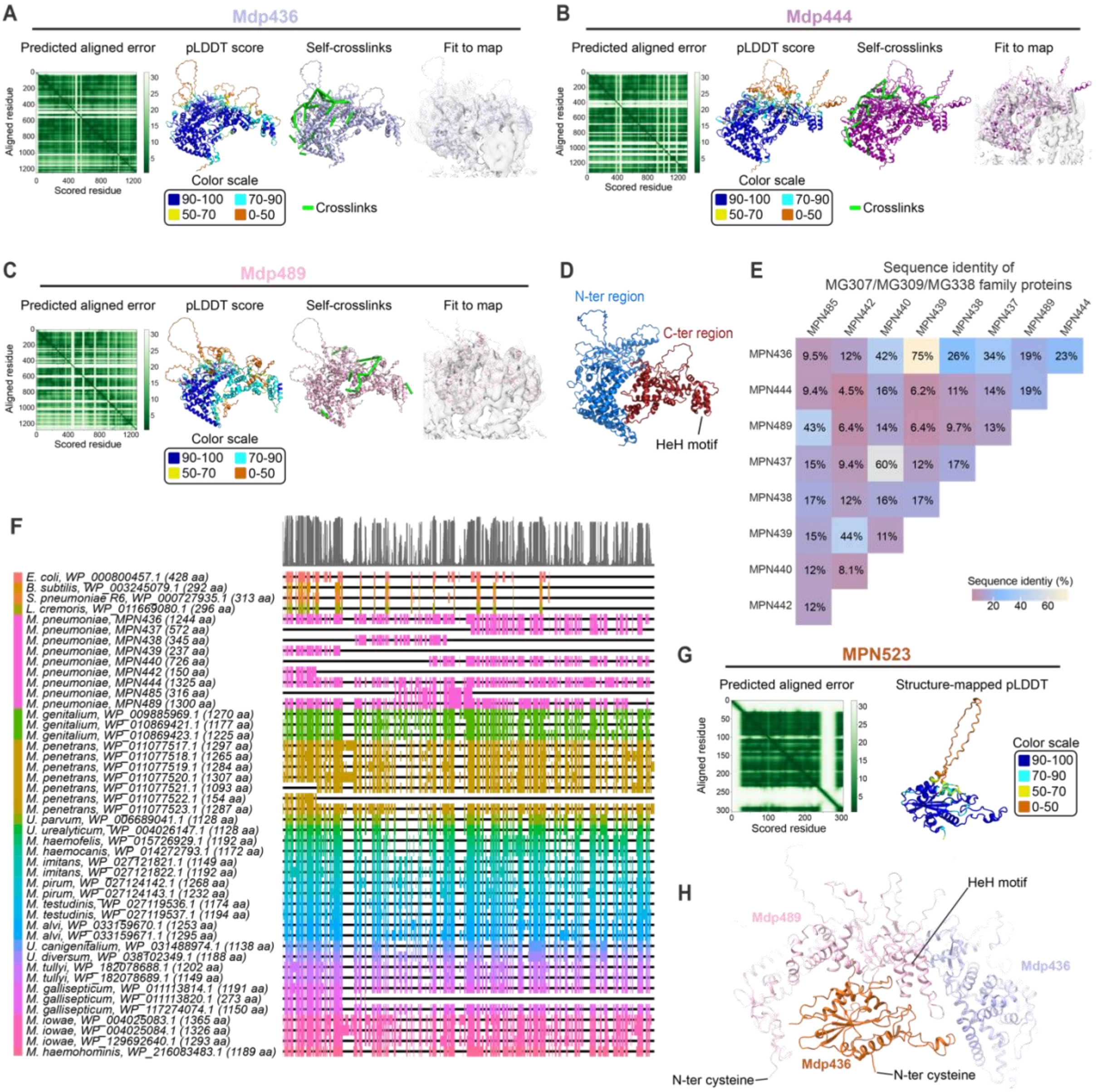


**Supplementary Figure 2. Details of the extracellular dome-complex proteins.** Related to Figure 2.

(A-C) AlphaFold2 predictions for the three Mdps and validation. From left to right: 2D distogram plot of predicted aligned error, pLDDT score mapped to the predicted model, the self-crosslinks in green mapped to the predicted model, and the best fit of model to map by rigid body fitting shown for (A) Mdp436, (B) Mdp444, (C) Mdp489. All self-crosslinks at a 5 % false discovery rate were satisfied by the AlphaFold2 predicted models. (D) The N-terminal region (blue) and the C-terminal region (red) of the AlphaFold model of Mdp436. (E) Heat-map showing the sequence identity between the MG307/MG309/MG338 family of proteins in *M. pneumoniae*. (F) Sequence alignment of proteins from the MG307/MG309/MG338 family and their homologs in other bacteria. The boxes indicate aligned residues and the black line indicates gaps in the alignment. The top grey histogram indicates conservation as a function of residue number. Different colored lines indicate different species. (G) AlphaFold2 prediction and validation of MPN523. The predicted aligned error is shown as a 2D distogram and the pLDDT scores are mapped on the predicted model. (H) Model of the interaction between Mdp436, Mdp489, and MPN523, showing that the HeH motif and C-ter region of Mdp489 interact with MPN523, and a long loop extends from MPN523 towards Mdp436.

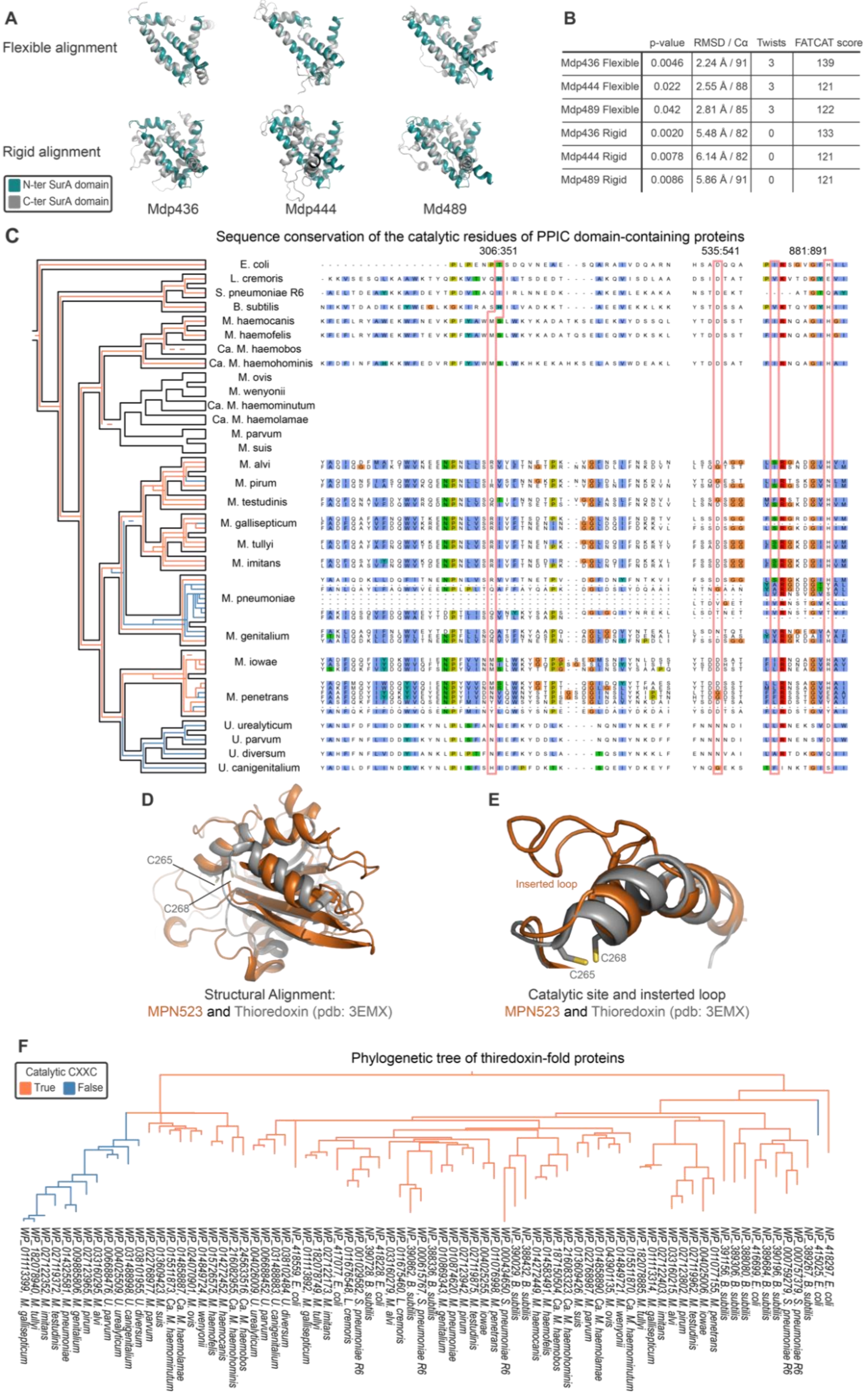


| | p-value | RMSD / Cα | Twists | FATCAT score |
|---|---|---|---|---|
| Mdp436 Flexible | 0.0046 | 2.24 Å / 91 | 3 | 139 |
| Mdp444 Flexible | 0.022 | 2.55 Å / 88 | 3 | 121 |
| Mdp489 Flexible | 0.042 | 2.81 Å / 85 | 3 | 122 |
| Mdp436 Rigid | 0.0020 | 5.48 Å / 82 | 0 | 133 |
| Mdp444 Rigid | 0.0078 | 6.14 Å / 82 | 0 | 121 |
| Mdp489 Rigid | 0.0086 | 5.86 Å / 91 | 0 | 121 |

**Supplementary Figure 3. Details of the bioinformatic analysis of the extracellular dome proteins.** Related to Figure 3.

(A) Alignment between the N- and C-terminal SurA domains in Mdp436, Mdp444, and Mdp489 using both rigid and flexible alignment mode of FATCAT. (B) Statistics for the different fits using FATCAT shown in A. All fits are significant, indicating that the regions are homologous. (C) Sequence conservation of the PPIC domain in the Mdps. Left: the phylogenetic tree is similar to the one shown in Figure 3E. Right: the sequence alignment of the PPIC domain in comparison to the PPIC domain of PrsA in the other species. The salmon-colored boxes indicate the position of the catalytic residues. The number on top of the alignment indicates the numbering in the consensus alignment. The alignment is colored according to the Clustal color scheme. For the phylogenetic tree, the black outline indicates the species tree, whereas the colored lines indicate the proteins: the orange-colored lines indicate that the catalytic residues in the PPIC domain are conserved, whereas the blue lines indicate that they are not conserved. The line color of a node is orange if at least one of its descendants has the catalytic residues. (D) Structural alignment of MPN523 and thioredoxin shows a conserved overall fold of the two proteins. The catalytic cysteines present in thioredoxin are indicated, and are not conserved in MPN523. (E) The substrate binding region of MPN523 and thioredoxin is conserved, but there is a large inserted loop in the helix of MPN523. (F) Phylogenetic tree of the proteins in the MPN523 family. The orange-colored lines indicate that the protein has retained the catalytic CXXC motif, whereas the blue lines indicate it is not retained.

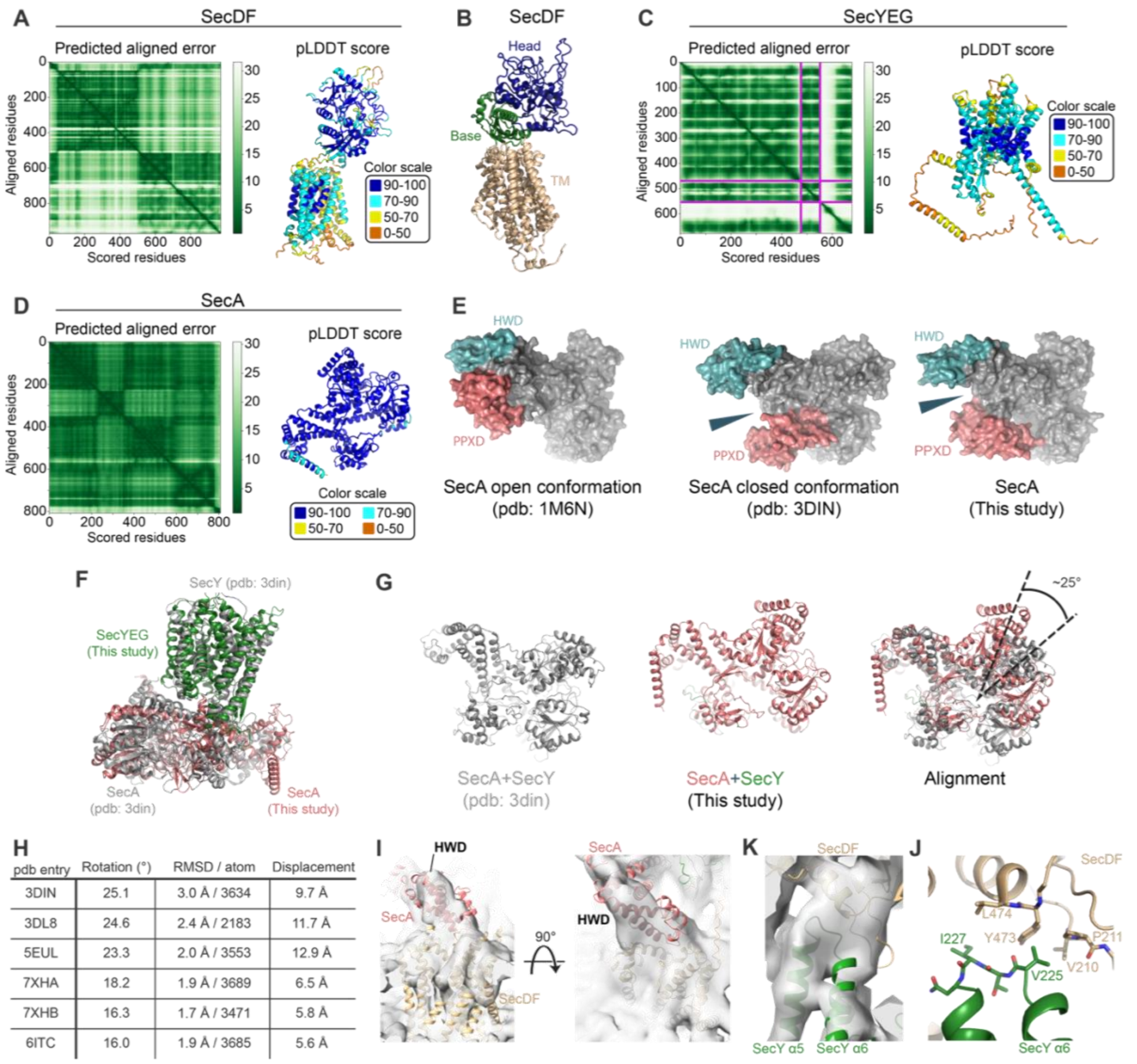


| pdb entry | Rotation (°) | RMSD / atom | Displacement |
|---|---|---|---|
| 3DIN | 25.1 | 3.0 Å / 3634 | 9.7 Å |
| 3DL8 | 24.6 | 2.4 Å / 2183 | 11.7 Å |
| 5EUL | 23.3 | 2.0 Å / 3553 | 12.9 Å |
| 7XHA | 18.2 | 1.9 Å / 3689 | 6.5 Å |
| 7XHB | 16.3 | 1.7 Å / 3471 | 5.8 Å |
| 6ITC | 16.0 | 1.9 Å / 3685 | 5.6 Å |

**Supplementary Figure 4. Details of the structural analysis of the Sec proteins.** Related to Figure 4.

(A) AlphaFold2 model and validation for the *M. pneumoniae* SecDF showing a 2D distogram of the predicted aligned error, and the pLDDT scores mapped to the predicted model. (B) The AlphaFold2 prediction of SecDF with the transmembrane (TM; beige), base (green), and head (blue) regions indicated. (C) Model and validation for the AlphaFold2 multimer prediction of the *M. pneumoniae* SecY, SecG, and SecE, showing the predicted aligned error plot and the pLDDT scores mapped to the model. The predicted aligned error is shown as a 2D distogram, where the first protein (from left to right; borders between proteins are indicated by magenta lines) corresponds to SecY, the second to SecG, and the third to SecE. (D) AlphaFold2 model and validation for the *M. pneumoniae* SecA showing a 2D distogram of the predicted aligned error, and the pLDDT scores mapped to the predicted model. (E) Comparison of the SecA structures in the open (pdb: 1M6N) and closed conformation (pdb: 3DIN), to the SecA major conformation derived in this study, showing that SecA is in the closed conformation. (F-G) Alignment of the SecY-SecA complex structure (pdb: 3DIN; grey) to structural model derived in this study (F). (G) Alignment of the SecA (pdb: 3DIN; grey) to the model in our study shows a relative rotation of 25°. (H) Table summarizing the rotation between the SecA-SecY subcomplex from our model in comparison to available *in vitro* structures, showing a range of 16-25°. (I) The interaction between SecA and SecDF shown in the context of the cryo-ET map of the closed conformation, showing that the SecA HWD interacts with the cytosolic tail of SecDF. (K) Detailed view of the SecY α5-α6 interaction with SecDF. (J) Potential interaction site between SecY and SecDF. Several hydrophobic residues likely to be involved in the interaction are shown.

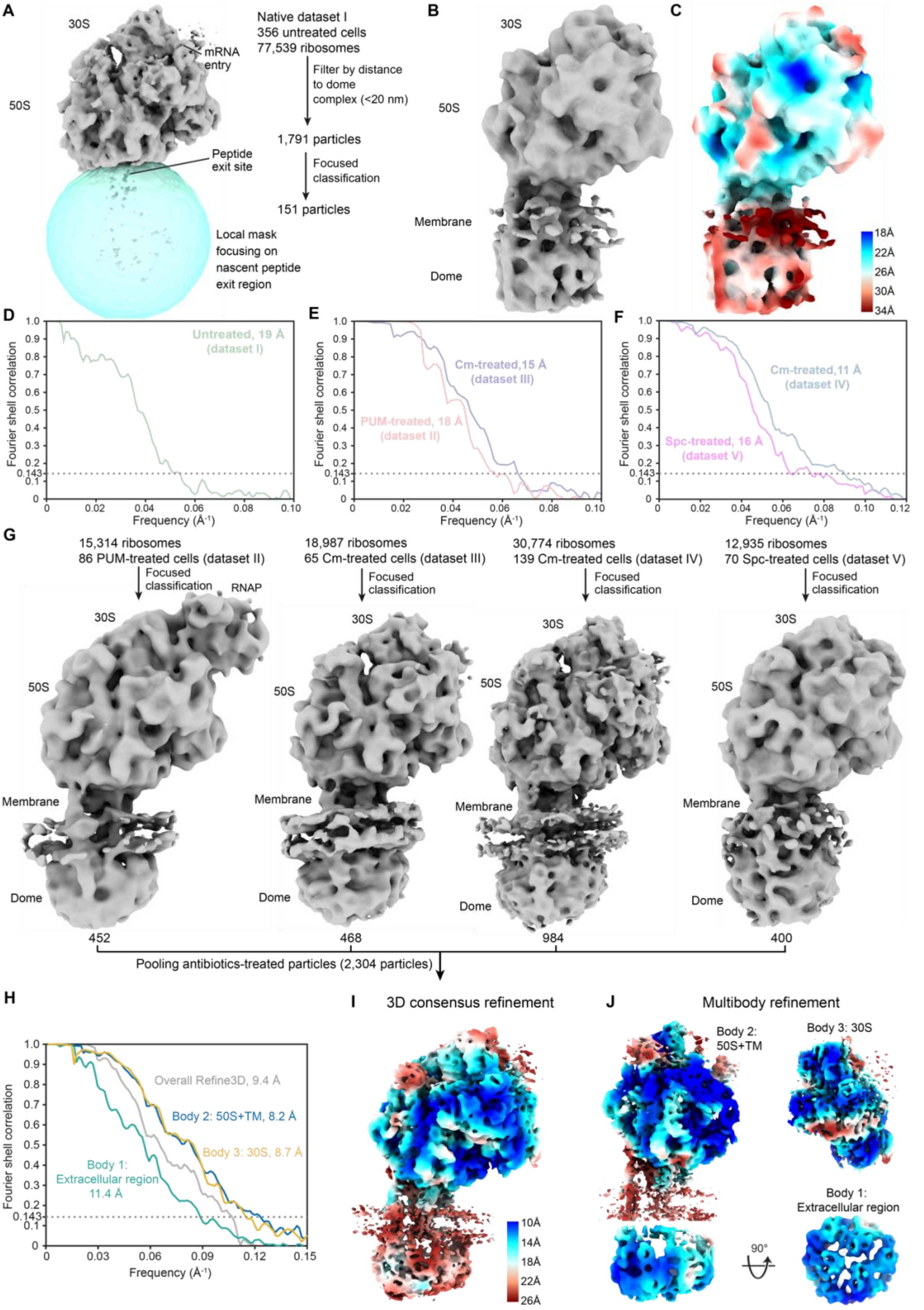


**Supplementary Figure 5. Cryo-ET data processing of the ribosome-dome complex.** Related to Figure 5.

(A) Classification of ribosome subtomograms with a spherical mask focused on the peptide exit site. To counter the high heterogeneity of ribosomes in native untreated cells, ribosomes were first sorted based on whether a dome complex was localized in close spatial proximity (20 nm cutoff). (B-D) Subtomogram average from the 151 classified ribosome-dome complexes in untreated cells, the corresponding local resolution map (C) and FSC curve (D). (G) Subtomogram analysis of the four antibiotic-treatment datasets. Particle numbers are indicated under the maps. RNAP: RNA polymerase stalled by PUM results in a physical collision with the ribosome. The classified complexes in the different antibiotic treatments exhibited similar overall architectures and were pooled for further refinement. (E-F) FSC curves for the ribosome-dome refinement in each of the antibiotic-treatment datasets. (H) FSC curves for global (overall Refine 3D) and multi-body refinement of the pooled ribosome-dome complexes in the antibiotic-treatment datasets. (I) Cryo-ET consensus map of the 2,304 pooled particles, colored by local resolution. (J) Multibody refinement results of the pooled particles, colored by local resolution.

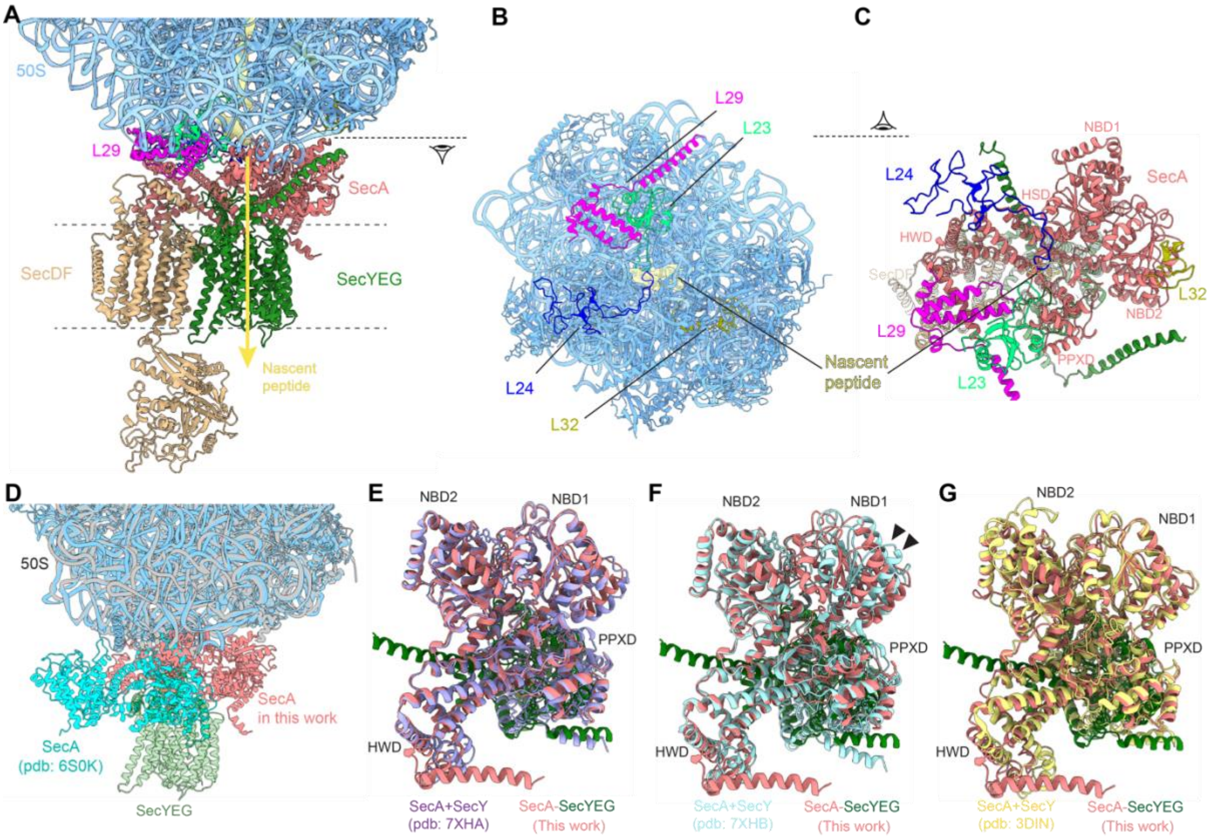


**Supplementary Figure 6. Modeling details of the ribosome-dome complex.** Related to Figure 5.

(A) Part of the model derived in this study for the ribosome-dome complex showing the interaction between the ribosome and the sec proteins. The putative path of the nascent peptide (density colored in yellow in the 50S) through SecA and the SecYEG channel is indicated by the yellow arrow. (B) View of ribosomal proteins L23, L24 and L29 at the interaction interface with SecA from the direction of SecA. The nascent peptide density inside the ribosome is shown in yellow. (C) View of the SecA interaction interface from the ribosome side. (D) Superposition of the *in vitro* ribosome-SecA complex (pdb: 6S0K; cyan and grey) and the ribosome-dome complex model derived in this study, aligned on the 50S. SecA is rotated approximately 180° between the two models. The *in vitro* structure does not allow simultaneous binding of SecYEG. (E-G) Comparison to previous SecA-SecY structures in the presence of a polypeptide substrate together with a nonhydrolyzable ATP analog (E: pdb: 7XHA) or ADP (F: pdb: 7XHB), or without a substrate (G; pdb: 3DIN). The ATP-bound state with substrate shows the highest structural similarity to the model derived in this study. Note the NBD1 domain (F: indicated by black arrowheads).

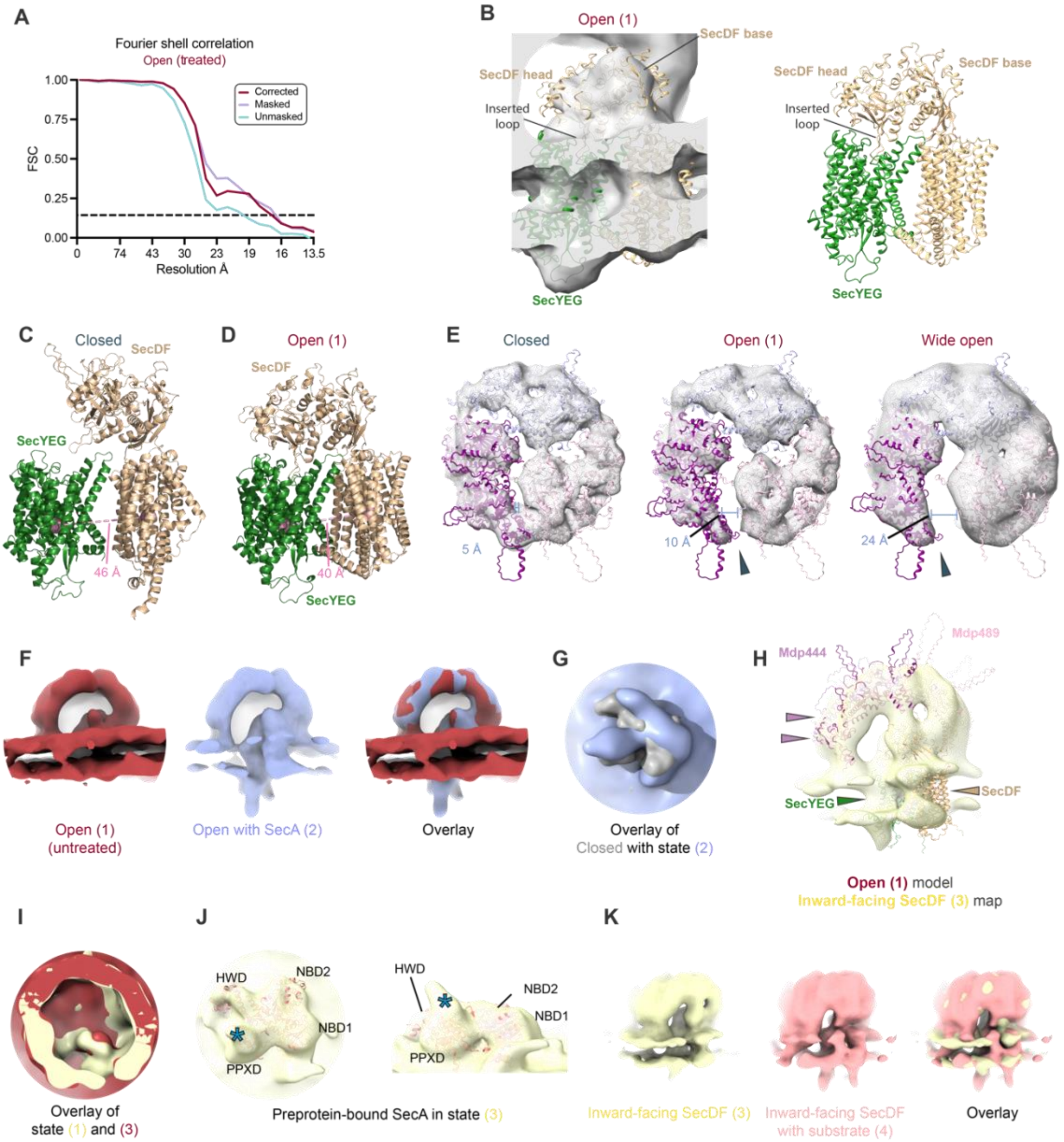


**Supplementary Figure 7. Details of the open and additional minor conformations of the dome complex.** Related to Figure 6.

(A) FSC plots for the open conformation of the dome complex pooled from antibiotics-treated cells. (B) SecYEG-SecDF multimer model from AlphaFold3 shown fitted in the cryo-ET map of the open conformation, showing that the model explains both the extracellular and membrane regions. Both the head and base region of SecDF fit the map, and the SecDF loop mediating SecDF-SecYEG interaction in the closed conformation is modeled inserted into the exit tunnel of SecYEG. (C-D) Comparison of the SecYEG-SecDF models from the closed (C) and open (D) conformation. The distance between the center of mass of SecDF and SecYEG decreases in the transition from the closed to the open conformation. (E) The maps and rigid-body fitted models of the Mdps shown for the closed, open and wide-open conformations of the complex. A large degree of flexibility in the Mdp444 and Mdp489 interface allows an opening of up to 24 Å. The distances between residue 178 in Mdp489 and residue 1124 in Mdp444 are indicated in each map. (F) Comparison of the open (1; red) and open with SecA (2; blue) conformations, showing the maps are almost identical except for the presence of the cytosolic SecA density in state 2. (G) Comparison of the closed (grey) and open with SecA (2; blue) conformations, showing that SecA is rotated between the two conformations. (H) The cryo-ET map of the inward-facing SecDF conformation (3; yellow) fitted with the structural model of the open conformation. Arrows indicate regions where Mdp444 (purple), SecYEG (green), and SecDF (beige) do not fit the map. (I) Comparison of the open (1; red) and inward-facing SecDF (3; yellow) conformations, showing a slight rotation of SecDF. (J) AlphaFold2 prediction of SecA fitted to the SecA density in the inward-facing SecDF state (3) shows that SecA is preprotein-bound. Preprotein density is indicated by a blue asterisk (*). (K) Comparison of the inward-facing SecDF (3; yellow) and inward-facing SecDF with substrate (4; pink) conformations, showing their similarity.

### Supplementary Table 1. Cryo-EM Data collection and image processing details.

| **Dome complex** | | | | | | | | |
|---|---|---|---|---|---|---|---|---|
| **Perturbation** | Native | | | | | | | PUM/ Cm |
| **Class** | Closed<br><br>SecA Major | Closed<br><br>SecA Minor | Open (1) no SecA | Open (2) with SecA | Intermediate (3) Inward-facing SecDF | Intermediate (4) Inward-facing SecDF with substrate | Mdp 444 Minor | Open (1) no SecA |
| **Magnification** | 81,000 | | | | | | | 81,000 /64,000 |
| **Voltage (kV)** | 300 | | | | | | | 300 |
| **Electron exposure $(e^{-1}/Å^2)$** | 115-135 | | | | | | | 115-137 |
| **Defocus range (µm)** | 1.5 to 3.5 | | | | | | | 1 to 3.5 |
| **Pixel size (Å/pixel)** | 1.7 | | | | | | | 1.7/1.3 |
| **Tomograms (no.)** | 355 | | | | | | | 183 |
| **Initial particle subtomograms (no.)** | 13,770 | 13,770 | 13,770 | 13,770 | 13,770 | 13,770 | 13,770 | 6,103 |
| **Final particle subtomograms (no.)** | 11,557 | 2,213 | 177 | 235 | 459 | 588 | 2,010 | 1,001 |
| **Map resolution (Å), FSC threshold 0.143** | 9.1 | 13 | 27 | 29 | 25 | 25 | 14 | 17 |

| **Ribosome-dome complex** | | | | | | | | |
|---|---|---|---|---|---|---|---|---|
| **Perturbation** | Native (I) | PUM (II) | Spectinomycin (V) | Cm (III) | Cm (IV) | Combined (II-V) | | |
| **Class** | Consensus refinement | | | | | Multibody Refinement | | |
| | | | | | | 30S | 50S+TM | Dome |
| **Magnification** | 81,000 | | | | 64,000 | 64,000-81,000 | | |
| **Voltage (kV)** | 300 | | | | 300 | 300 | | |
| **Electron exposure $(e^{-1}/Å^2)$** | 115-137 | | | | 137 | 115-137 | | |
| **Defocus range (µm)** | 1.5 to 3.5 | | | | 1 to 3.25 | 1 to 3.25 | | |
| **Pixel size (Å/pixel)** | 1.7 | | | | 1.3 | 1.3-1.7 | | |
| **Tomograms (no.)** | 355 | 86 | 70 | 65 | 139 | 716 | | |
| **Initial particle subtomograms (no.)** | 77,539 | 15,314 | 13,935 | 18,987 | 30,774 | 2,304 | 2,304 | 2,304 |
| **Final particle subtomograms (no.)** | 151 | 452 | 400 | 468 | 984 | 2,304 | 2,304 | 2,304 |
| **Map resolution (Å), FSC threshold 0.143** | 19 | 18 | 16 | 15 | 11 | 8.7 | 8.2 | 11.4 |

**Supplementary Table 2. Surface shaving mass spectrometry.** Quantified fold change at the peptide level between treated (trypsin or proteinase K) or untreated *M. pneumoniae* cells. Related to Figure 2A.

**Supplementary Table 3. Rigid-body refinement into the dome complex map using PowerFit.** Cross-correlation scores for the fitting of the AlphaFold prediction for each candidate protein to the extracellular region of the dome complex map in the closed conformation. The proteins are indicated by their gene name. Related to Figure 2B.

**Supplementary Table 4. Output of the FoldSeek domain search.** Raw output of the FoldSeek search for the Mdps and MPN523, listing the hits for each domain searched, for both TM_align and 3DI/AA. Related to Figure 3B.

**Supplementary Table 5. Output of the DALI domain search.** Raw output of the DALI search for the Mdps and MPN523, listing the hits for each domain searched. Related to Figure 3B.
